# Single-cell multi-omics maps clonal IEL expansion and epithelial remodelling in refractory coeliac disease

**DOI:** 10.64898/2026.08.30.747985

**Authors:** Michael Li, Arman Safavi, Jerome Samir, Raymond Louie, Esmaeil Roohparvar Basmenj, Martina Bonomi, Thiruni Adikari, Brian Gloss, Jun Xing, Katherine Jackson, Andrew Calcino, Dan Suan, Valentina Vaira, Claire Milthorpe, Melinda Hardy, Etienne Masle-Farquhar, Scott Read, Matt Field, Luca Elli, Marco Vincenzo Lenti, Antonio Di Sabatino, Rachele Ciccocioppo, Jason A. Tye-Din, Golo Ahlenstiel, Christopher C. Goodnow, Mandeep Singh, Fabio Luciani

**Affiliations:** University of New South Wales, Kensington NSW, Sydney, Australia; The Westmead Institute for Medical Research, Westmead NSW, Sydney, Australia; Mutinex, Sydney, Australia; University of Oxford, Department of Paediatrics; Garvan Institute of Medical Research, Darlinghurst NSW, Sydney, Australia; James Cook University, Townsville, QLD, Australia; Western Sydney University, Western Sydney Local Health District, Sydney NSW, Australia; University of Milan, Milan, Italy; Department of Internal Medicine and Therapeutics, University of Pavia, Fondazione IRCCS Policlinico San Matteo, Pavia, Italy; University of Chieti, Gastroenterology & Endoscopy Unit, Department of Medicine and Ageing, Italy; Walter and Eliza Hall Institute of Medical Research (WEHI), Victoria, Australia

## Abstract

**Background:** Refractory coeliac disease type 1 (RCD1) lacks defining molecular markers, and the immune and epithelial mechanisms sustaining intestinal injury remain poorly understood.

**Objective:** To define the clonal immune and epithelial states that distinguish RCD1 from active coeliac disease (ACD) and determine their spatial organisation in the duodenal mucosa.

**Design:** We integrated single-cell RNA sequencing, CITE-seq surface proteomics and paired T-cell receptor sequencing of duodenal immune and epithelial compartments from Healthy controls (n = 6), ACD (n = 7), RCD1 (n = 9) and RCD2 (n = 2), with spatial transcriptomics in a subset of biopsies.

**Results:** RCD1 showed widespread TCRαβ and TCRγδ clonal expansion across multiple IEL states, extending beyond previously defined mutation-bearing aberrant clones. Distinct IEL populations converged on a shared programme of adaptive persistence, innate-like signalling, metabolic fitness and cytoskeletal remodelling; *GZMK* expression marked both clonally expanded and non-clonal disease-associated states. In parallel, RCD1 epithelium showed loss of mature absorptive cell states and expansion of stress-associated, immune-interacting and regenerative programmes. Transit-amplifying cells acquired differentiation and tissue-remodelling signatures, while enteroendocrine cells expanded and developed a sensory-neurosecretory programme involving *TRPA1, TRPV1*, vesicle trafficking and *NEUROD1* regulon activity. Spatial transcriptomics localised regenerative and enteroendocrine-associated epithelial programmes adjacent to immune-visible epithelial regions and *KLRK1*/*GZMK*-expressing IEL-rich niches in refractory tissue.

**Conclusion:** RCD1 represents a distinct mucosal state characterised by coordinated clonal IEL adaptation and epithelial remodelling, rather than simple amplification of ACD, providing a cellular framework for persistent tissue injury and disease stratification.

## Introduction

Coeliac disease (CeD) is a chronic immune-mediated enteropathy triggered by dietary gluten in genetically susceptible individuals, affecting approximately 1% of the global population [1]. Although most patients improve following a strict gluten-free diet (GFD), up to 30% continue to experience persistent symptoms, mucosal inflammation and villous abnormalities despite apparent dietary adherence [2, 3]. Furthermore, follow-up biopsy studies have further demonstrated that villous atrophy frequently persists even in patients considered clinically well controlled on a GFD [4, 5, 6], suggesting that ongoing tissue remodelling and immune activation may continue despite treatment.

A small subset of patients, estimated at approximately 1% of all CeD cases, develop refractory coeliac disease (RCD), a severe condition characterised by persistent malabsorption, villous atrophy and symptoms despite more than 12 months on a strict GFD [2]. RCD is associated with significant morbidity and an increased risk of life-threatening complications, including enteropathy-associated T-cell lymphoma (EATL) [7, 8]. Traditionally, RCD has been divided into two subtypes. RCD type 2 (RCD2) is characterised by the expansion of aberrant intraepithelial lymphocytes (IELs), typically defined by loss of surface (s)CD3 and T cell receptor (TCR) expression, retention of intracellular CD3ε, loss of CD8, expression of markers such as CD7 and CD103, and clonal TCRγ gene rearrangement [9, 10, 11]. The presence of more than 20% aberrant IELs by flow cytometry is a key diagnostic criterion and identifies patients at a particularly high risk of progression to lymphoma [12]. By contrast, RCD type 1 (RCD1) lacks a defining aberrant IEL population and is generally diagnosed by exclusion after ruling out ongoing gluten exposure, alternative enteropathies, and overt malignancy [2].

The molecular basis of RCD2 has been extensively investigated over the past decade. Somatic mutations affecting lymphoma-associated pathways, particularly the JAK-STAT signalling axis, are frequently detected within aberrant IEL populations [13, 14]. These alterations confer gain-of-function signalling, enhancing responsiveness to IL-15 and promoting aberrant IEL survival and expansion [15, 16]. Additional recurrent alterations include mutations in epigenetic regulators such as *TET2* and *KMT2D* and the RNA helicase *DDX3X*, as well as structural and chromosomal abnormalities including recurrent gains of chromosome 9q and loss of heterozygosity at the *JAK1* locus [14, 17, 18]. These clones are directly implicated in the pathogenesis of villous atrophy, as they express high levels of NK-associated activation and cytotoxic genes and have been shown to directly kill intestinal epithelial cells *in vitro* [19, 20, 21]. The cellular origin of aberrant IELs has been linked to a rare population of IL-15-dependent innate-like lymphocytes that exhibit both NK-cell and T-cell characteristics and reside within the normal intestinal epithelium [11, 15, 22]. Together, these findings have established RCD2 as a clonal lymphoproliferative disorder arising within the intestinal immune compartment.

Beyond serving as a target of immune-mediated injury, the intestinal epithelium undergoes extensive functional and developmental remodelling in active coeliac disease (ACD) [23, 24]. Recent single-cell studies have demonstrated a shift from mature absorptive enterocytes towards proliferating stem, transit-amplifying and early enterocyte states, accompanied by impaired lipid, carbohydrate, vitamin and mineral transport programmes and broad interferon-driven induction of antigen-presentation and inflammatory pathways [23, 24]. Secretory lineages are also altered, with increased secretory epithelial populations and changes in chemokine, antimicrobial and gut-hormone expression. Notably, FitzPatrick *et al.* identified perturbation of the enteroendocrine compartment, including expansion of *NEUROG3*-positive endocrine progenitors and somatostatin-producing D cells, highlighting disruption of the sensory and neuroendocrine functions of the duodenal epithelium. Some progenitor and enteroendocrine abnormalities persisted following GFD, suggesting a lasting immune–epithelial dysregulation. Richards *et al.* further implicated coordinated T-cell– myeloid–stromal–epithelial signalling in this remodelling, whereby lymphoid-derived IFN-γ and myeloid-derived IL-1β reprogram fibroblasts to support crypt expansion and epithelial differentiation. However, whether these epithelial changes intensify or diversify in RCD remains largely unknown.

These epithelial knowledge gaps parallel the continuing uncertainty surrounding the immune mechanisms that sustain intestinal inflammation in RCD1. Patients with RCD1 generally lack large expansions of canonical aberrant IELs and often present with heterogeneous clinical manifestations, making diagnosis challenging [2, 25]. Recent genomic studies, however, have challenged the traditional distinction between RCD1 and RCD2. We and others have recently demonstrated that clonally expanded IEL populations carrying somatic mutations in lymphoma driver genes, particularly within the JAK/STAT pathway, can also be detected in RCD1. While some of these clones display classical RCD2-like phenotypes, the majority are found within conventional CD8^+^ T-cell populations retaining sCD3 and TCR expression [18, 26]. These findings reveal a previously unappreciated heterogeneity of aberrant clones in RCD1 and suggest that clonal expansion and somatic evolution may represent shared pathogenic mechanisms across the spectrum of refractory disease. While RCD is marked by clonal expansions of IELs carrying somatic driver mutations, considerably less is known about whether broader immune remodelling occurs across other compartments, including non-mutant IEL populations, lamina propria T-cells, and B-cells, or whether the epithelial compartment undergoes disease-specific changes. IELs play a central role in tissue destruction in CeD, with gliadin-induced IL-15 production by enterocytes and subepithelial myeloid cells promoting upregulation of stress ligands including MICA, MICB and HLA-E on the enterocyte surface, and thereby triggering NK-receptor-mediated killing by both TCRαβ and TCRγδ IEL populations [19, 27, 28]. Recent single-cell RNA sequencing studies have begun to resolve the cellular basis of these changes in ACD, demonstrating interferon-driven transcriptional reprogramming, a shift towards epithelial progenitor states, and loss of absorptive enterocytes, consistent with crypt hyperplasia and villous atrophy [23, 24]. However, it remains unknown whether these epithelial changes represent a passive consequence of immune-mediated tissue damage or an active component of disease pathogenesis. It is also unclear how the epithelial and broader immune landscape is further remodelled in refractory disease.

These observations raise several fundamental questions: are RCD1 and RCD2 distinct biological entities or different manifestations of a shared disease process; how are aberrant clonal expansions distributed across intestinal immune populations; and how does chronic inflammation reshape epithelial differentiation programmes in refractory disease? To address these questions, we performed single-cell multi-omics profiling of small intestinal biopsies from healthy controls and patients with ACD, RCD1 and RCD2. By integrating transcriptomic, surface protein and TCR information, we generated a comprehensive map of immune and epithelial cell states across the spectrum of diseases. We demonstrate that RCD1 retains several features already evident in ACD, including inflammatory activation, clonal expansion of IEL populations and selected stress-associated transcriptional programmes, while also acquiring molecular features that distinguish it from ACD. These include recurrent adaptive remodelling across multiple IEL states and distinct epithelial programmes, with changes emerging at the progenitor stage and extending to specialised regenerative, metabolic and sensory epithelial states.

## Results

To identify novel molecular signatures underlying RCD1, we performed single-cell RNA sequencing (scRNA-seq) of paired immune and epithelial compartments from small intestinal biopsies obtained from healthy controls (*n* = 6), ACD (*n* = 7), RCD1 (*n* = 9) and RCD2 (*n* = 2), comprising a total of 24 individuals **(Fig. 1A, Supplementary Table 1)**. Immune (CD45) and epithelial (EPCAM) cells were isolated in parallel by flow cytometric sorting **(Supplementary Fig. 1A)** and profiled using the 10x Genomics platform. Immune cells were additionally subjected to paired TCR sequencing [29] and quantification of up to 155 surface proteins using CITE-seq [30] **(Fig. 1A, Supplementary Table 2)**. Following quality control, a total of 162,630 immune cells and 26,230 epithelial cells were analysed.

**Figure 1.**
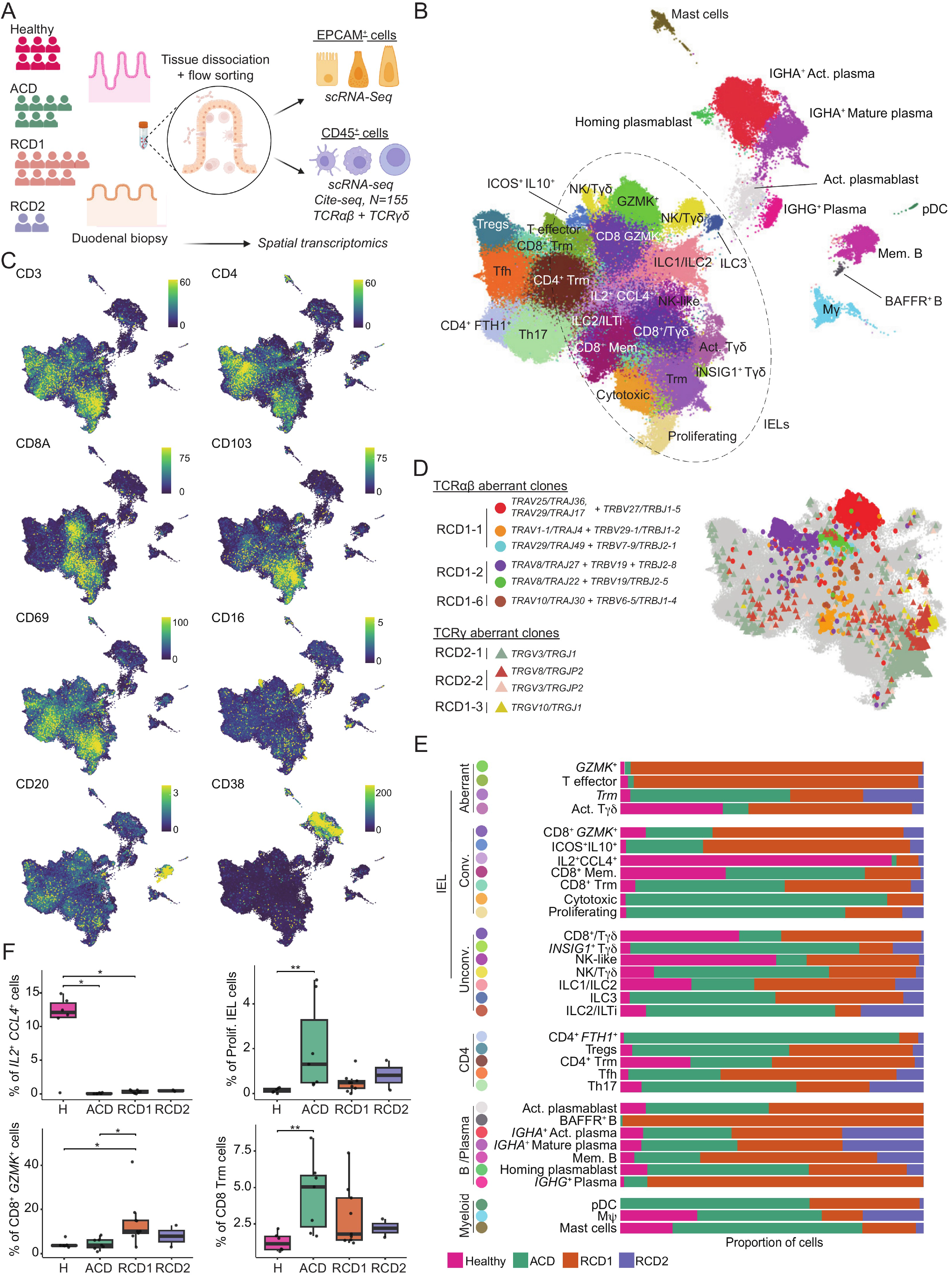
The immune cells from small intestine of healthy individuals and coeliac patients showed diverse immunophenotypic profile. **A)** Schematic representation of the study plan, number of patients enrolled, data collection and single-cell experiments in this study. **B)** UMAP representation of the immune cells obtained from small intestine biopsies of 24 individuals (Ncells = 162630). **C)** UMAP representations show the normalised expression levels of a selective group of protein markers. **D)** The distribution of previously identified aberrant clones across UMAP. **E)** Stacked bar plots show the relative contribution of cells from Healthy, ACD, RCD1 and RCD2 samples to each immune cell cluster. **F)** The box plots show the percentage of cells retrieved from each disease state for specific clusters. Stars represent statistical significancy calculated using two-sided Wilcoxon rank-sum tests. Boxes show the median and interquartile range, and whiskers extend to 1.5 times the interquartile range. ACD: Active Coeliac Disease, RCD1: Refractory Coeliac Disease Type 1, RCD2: Refractory Coeliac Disease Type 2, scRNA seq: single-cell RNA sequencing, Trm: T resident memory, Act. Tγδ: Activated γδ T cells, CD8^+^ Mem.: CD8^+^ memory T cells, ILC: Innate lymphocyte cells, Act. Plasmablast: Activated plasmablast, IGHA^+^ Act. Plasma: IGHA^+^ Activated plasma cells, pDC: Plasmacytoid dendritic cell

Analysis of the immune compartment identified 33 transcriptionally distinct populations, comprising 16 conventional T-cell clusters defined by rearranged TCR β chains, 7 unconventional immune clusters, encompassing NK cells, innate lymphoid cells (ILCs), and γδ T-cells, 7 B-cell/plasma cell clusters, and 3 myeloid populations, with cells from individual patients distributed across the integrated UMAP **(Fig. 1B, Supplementary Fig. 1B-C)**. Cell type annotations were further supported by canonical surface protein markers (**Fig. 1C, Supplementary Fig. 1D**). IELs were identified by surface expression of CD103, a characteristic marker of intestinal IELs [31, 32], and comprised the majority of immune cells (**Fig. 1B, Supplementary Fig. 1B**). IEL populations comprised conventional TCRαβ subsets together with multiple γδ T-cell and ILC1, 2, and 3. Conventional TCRαβ IELs were distributed across transcriptionally distinct clusters including cytotoxic, tissue-resident memory (Trm), proliferative and effector cell states **(Fig. 1B, Supplementary Fig. 1E)**, highlighting the marked heterogeneity of the IEL compartment.

### Aberrant lymphocytes occupy transcriptionally distinct immune states within the intestinal immune landscape

In our previous study, we identified expanded IEL clones carrying somatic driver mutations in both RCD1 and RCD2 [18]. These comprised aberrant TCRαβ clones with sCD3^+^ CD103^+^ phenotype and aberrant sCD3^−^ CD103^+^ clones identified through clonotypic TCRγ rearrangements. Our scRNA-seq dataset included 4 RCD1 and 2 RCD2 biopsies containing these expanded mutation-bearing IEL clones carrying somatic driver mutations (hereafter referred to as “aberrant clones”), enabling their transcriptional states to be resolved by integrating TCR sequences with the single-cell transcriptomic data [18]. sCD3^+^ TCRαβ aberrant clones were distributed across distinct transcriptional states, with the largest clones accumulating in the *GZMK*^+^ and T effector clusters **(Fig. 1D)**, characterised by expression of *GZMK* and *CD27* in the former and *CD40LG* and *RORA* in the latter **(Supplementary Fig. 1E, Supplementary Table 3)**. In contrast, sCD3^−^ TCRγ rearranged aberrant clones from RCD1 and RCD2 patients localised predominantly within the Trm and activated Tγδ clusters, characterised by higher expression of *GZMA* and *KIR2DL4* **(Supplementary Fig. 1E, Supplementary Table 3)**. The Trm cluster consisted of cells from all disease states and expressed higher levels of CD7, CD69 and CD103 proteins, together with NK-associated markers including NKp46, *KLRC2* (NKG2C), *KLRD1* (CD94) and *KLRK1* (NKG2D) **(Fig. 1C, Supplementary Fig. 1D, E, Supplementary Table 3)**.

The distribution of non-aberrant immune populations also changed markedly across disease states **(Fig. 1E, F, Supplementary Fig. 1B-C).** Healthy tissues showed a higher representation of *IL2^+^ CCL4^+^* IELs, which were markedly depleted in ACD (as previously shown [24] and remained infrequent in RCD. In contrast, ACD was associated with increased proportions of cytotoxic and proliferating IELs, and CD8^+^ Trm cells, consistent with the activation and proliferation of tissue-resident IELs during intestinal response to gluten [33]. Further remodelling was observed in RCD1, with a significant increase in the proportion of *GZMK*^+^ IELs. Together, these findings indicate that RCD1 is characterised not only by expansion of aberrant clones, but also by broader remodelling of the IELs immune compartment.

### Clonal expansion is broadly increased across **αβ** and **γδ** IEL populations in RCD1

To determine how T-cell clonal architecture differs across disease states, we integrated paired chain TCRαβ and TCRγδ repertoires with the transcriptionally defined T-cell populations **(Fig. 2A)**. T cells assigned TCRαβ and TCRγδ chains were distributed across multiple IEL states, whereas previously identified aberrant TCRαβ and TCRγ clones occupied a more restricted subset of these populations, with aberrant TCRαβ cells particularly enriched within *GZMK*^+^, T effector and related IEL states and aberrant TCRγ cells predominantly localising within γδ-containing populations (**Fig. 2A**). Analysis of clonotype size demonstrated that increased clonal expansion in RCD1 extended well beyond cell clusters associated with aberrant clones. Within the TCRαβ repertoire, large clonotypes were more prominent in RCD1 than healthy and ACD across multiple IEL populations, with increased clonal expansion observed across *GZMK*^+^, T effector, CD8^+^*GZMK*^+^, CD8^+^ memory and NK-like TCRαβ-containing cells **(Fig. 2B)**. Sample-level analysis supported increased clonal expansion within the TCRαβ compartment across all IELs, showing a reduction in the frequency of singleton TCRαβ IELs and an increased frequency of large TCRαβ clones in RCD1 (**Fig. 2C**). Consistent with this shift towards clonal dominance, repertoire diversity was reduced in RCD1 as measured by Shannon entropy (**Supplementary Fig. 2A)**.

**Figure 2.**
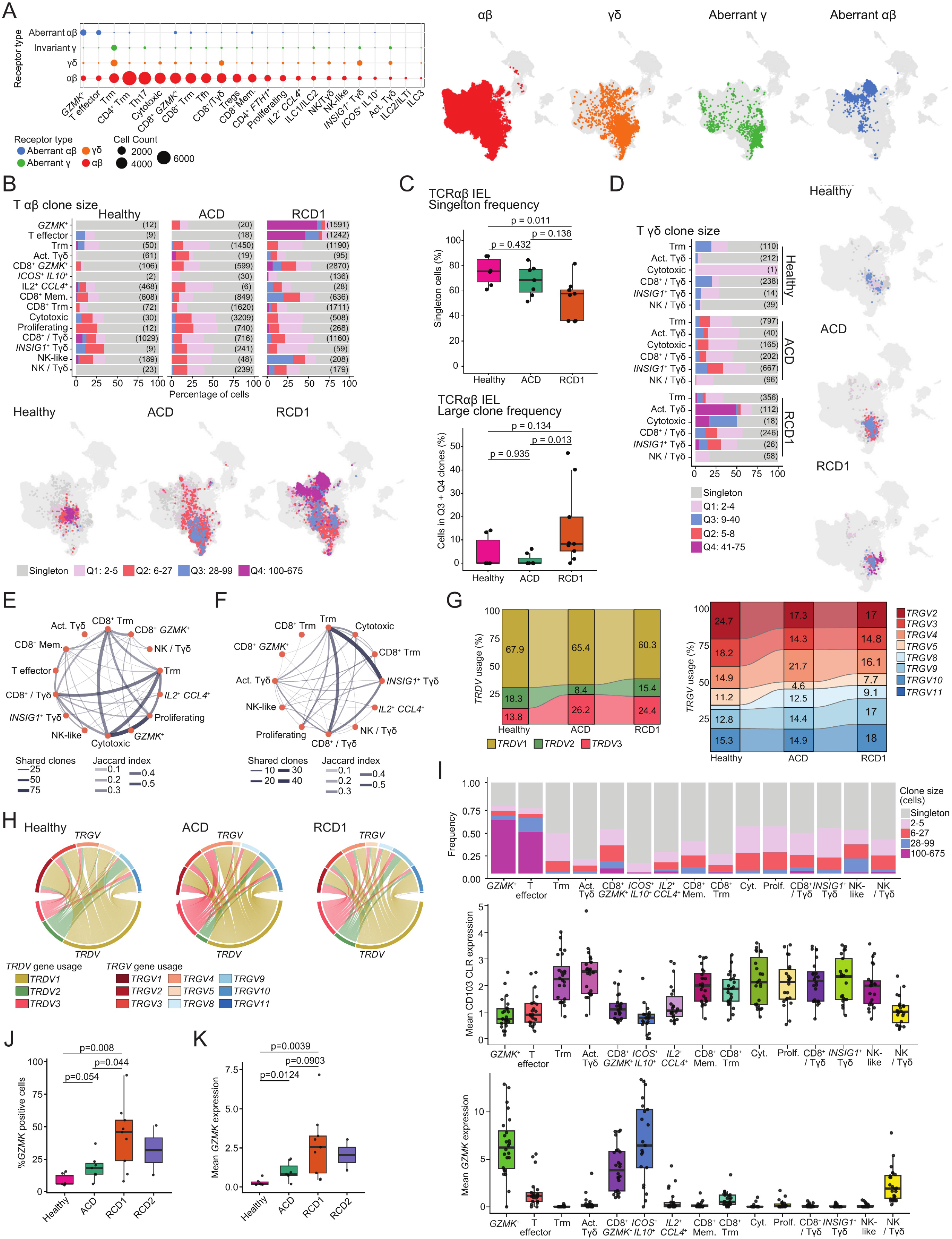
Clonally expanded IELs increase with disease severity. **A)** The dot plot shows the distribution of αβ, γδ, aberrant αβ and aberrant γδ receptors across T-cell clusters. Dot size represents the number of cells and colour indicates receptor type. UMAPs highlight cells of each receptor class in colour against all other cells in grey. **B)** Stacked bar plots represent the percentage of TCRαβ cells in each clone size category across IEL clusters (excluding ILC clusters) in Healthy, ACD and RCD1 samples. RCD2 samples are excluded from this analysis due to limited sample size. Only cells containing paired TCRαβ CDR3 sequences are included. Expanded clonotypes were divided into Q1-Q4 using the 25th, 50th and 75th percentiles of the pooled TCRαβ clone size distribution after excluding RCD2 samples. Values in parentheses indicate the total number of eligible cells in each cluster and diagnostic group. UMAPs below show the distribution of the same clone size categories across UMAP for each disease state, with all other cells shown in grey. **C)** Stacked bar plots show the corresponding TCRγδ clone size distributions across 6 IEL clusters containing the highest number of Tγδ cells in Healthy, ACD and RCD1 samples. RCD2 samples are excluded from this analysis due to limited sample size. Only cells containing paired TCRγδ CDR3 sequences are included. Expanded clonotypes were divided into Q1-Q4 using the 25th, 50th and 75th percentiles of the pooled TCRγδ clone size distribution after excluding RCD2 samples. Values in parentheses indicate the total number of eligible cells in each cluster and diagnostic group. UMAPs show the diagnosis-specific distribution of these clone size categories. **D)** Boxplots summarise TCRαβ clonality across IEL clusters at sample level. The top plot shows the percentage of singletons, and the bottom plot shows the percentage of cells assigned to large Q3 or Q4 clones, representing the upper half of the expanded clone distribution. For each sample, the number of cells containing paired TCRαβ CDR3 sequences was summed across IEL clusters, and each frequency was calculated by dividing the corresponding cell count by the total number of eligible cells. Each point represents one sample. Displayed *p*-values are from two-sided Wilcoxon rank-sum tests. **E-F)** Circular networks show the strongest intercluster shared-clonotype relationships for **E)** TCRαβ and **F)** TCRγδ across IEL clusters (excluding ILC clusters). Edge width represents the number of shared clonotypes, and edge opacity represents the Jaccard index between cluster-specific clonotype sets. **G)** Alluvial plots show percentage of *TRDV* and *TRGV* gene usage in Healthy, ACD and RCD1 samples among cells containing paired TCRγδ CDR3 sequences. **H)** Chord diagrams show *TRDV*-*TRGV* pairing patterns among cells containing paired TCRγδ CDR3 sequences in Healthy, ACD and RCD1 samples. **I)** Boxplots show sample-level mean normalised CD103 protein expression and sample-level mean *GZMK* gene expression across IEL clusters. Each point represents one sample-cluster summary. Boxes show the median and interquartile range, and whiskers extend to 1.5 times the interquartile range. **J-K)** Boxplots show sample-level **J)** percentage of GZMK-positive IELs and **K)** mean normalised *GZMK* RNA expression by diagnosis. GZMK positivity was defined as normalised RNA expression greater than zero. Each point represents one sample. Displayed *p*-values compare Healthy, ACD and RCD1 using two-sided Wilcoxon rank-sum tests. RCD2 is shown descriptively and was not included in the comparisons due to limited sample size. Boxes show the median and interquartile range, and whiskers extend to 1.5 times the interquartile range. ACD: active coeliac disease, IEL: intraepithelial lymphocyte, ILC: innate lymphoid cell, RCD1 and RCD2: refractory coeliac disease types 1 and 2

Expanded clonotypes were also observed within the TCRγδ repertoire, particularly among activated, cytotoxic and tissue-resident γδ-containing IEL states (**Fig. 2D**). However, TCRγδ clonality showed substantial inter-individual variability, and sample-level analysis did not identify a significant overall difference in singleton or large-clone frequencies between Healthy, ACD and RCD1 groups (**Supplementary Fig. 2B**). Thus, prominent γδ clonal expansion was observed in a subset of RCD1 patients but was not a consistent cohort-wide feature.Increased clonality was also detected among ILC and non-IEL CD4 T-cell populations in RCD1 compared with ACD and healthy controls **(Supplementary Fig. 2C, D)**. Together, these findings indicate that RCD1 is characterised by broad remodelling of the intestinal T-cell repertoire, with increased clonal dominance extending across both conventional and unconventional IEL populations rather than being restricted to mutation-bearing aberrant clones.

Clonal sharing analysis further demonstrated that individual expanded clonotypes occupy multiple transcriptionally distinct IEL states. Among TCRαβ cells, extensive clonotype sharing was observed across cytotoxic and proliferating populations and among CD8^+^/Tγδ, Trm and CD8^+^ Trm populations **(Fig. 2E)**. These findings indicate that clonally related IELs are not confined to a single transcriptional state, consistent with substantial phenotypic diversity within expanded clones.

Previous studies showed that γδ T-cell repertoires are markedly remodelled in ACD, where gluten-driven inflammation alters their abundance, phenotype and repertoire structure [34]. In our dataset, we analysed 4,051 Tγδ cells with paired γ- and δ-chain CDR3 sequences, together with 1,495 previously identified aberrant cells carrying clonotypic TCRγ rearrangements. These cells were distributed across multiple transcriptionally distinct IEL populations **(Fig. 2A)**, with the largest numbers found within the Trm, *INSIG1*^+^ Tγδ and CD8^+^/Tγδ clusters. The Trm population also contained the majority of aberrant TCRγ clones. TCRγδ clonotypes similarly displayed substantial inter-cluster sharing, particularly among Trm, *INSIG1*^+^ Tγδ and CD8^+^/Tγδ populations **(Fig. 2F)**. UpSet analysis further demonstrated sharing of individual TCRγδ clonotypes across multiple γδ IEL populations **(Supplementary Fig. 2F)**. The TCRγδ repertoire also displayed disease-associated changes in V-gene composition. *TRDV1* remained the predominant δ-chain gene across Healthy, ACD and RCD1 samples, consistent with the established enrichment of Vδ1^+^ γδ T cells within the human intestinal compartment [33, 34]. However, the relative contributions of TRDV2 and TRDV3, together with the distribution of TRGV genes, varied across disease states, indicating broader restructuring of the γδ repertoire (**Fig. 2G**). Analysis of paired *TRDV*– *TRGV* gene usage further revealed changes in γδ-chain pairing across Healthy, ACD and RCD1 samples (**Fig. 2H**). Whereas previous studies have described depletion of the physiological Vγ4δ1+ (*TRDV1*-*TRGV4* γδ T cell) intestinal IEL compartment in coeliac disease [33, 34], our data showed no significant depletion of this γδ T cell subset **(Supplementary Fig. 2G)**, thus highlighting the need for larger TCRγδ repertoires in clinical studies. Our data indicate that RCD1 is characterised by broader remodelling of the γδ compartment, involving altered clonal architecture and receptor composition across multiple γδ IEL states **(Fig. 2F–H; Supplementary Fig. 2F, G)**.

Together, these analyses demonstrate extensive remodelling of both αβ and γδ IEL repertoires in RCD1, characterised by increased clonal expansion, sharing of clonotypes across transcriptionally distinct IEL states, and altered γδ receptor composition. Notably, several of the populations containing prominent expanded αβ clonotypes were characterised by high *GZMK* expression, prompting us to examine whether GZMK represented a feature specific to aberrant clones or a broader disease-associated IEL state.

### *GZMK* expression marks distinct clonally expanded and non-clonal IEL states and increases in refractory disease

Our previous findings revealed increased expression of GZMK within mutation-bearing aberrant clones [18], and emerging evidence implicate GZMK-expressing T cells in chronically inflamed tissues and autoimmune disease [35, 36, 37]. We therefore asked whether increased *GZMK* expression in RCD simply reflected expansion of aberrant clones or represented a broader feature of the intestinal immune response. Analysis of *GZMK* expression across IEL populations identified several transcriptionally distinct populations with high expression, including *GZMK*^+^ IELs, CD8^+^*GZMK*^+^, *ICOS*^+^*IL10*^+^ and NK/Tγδ clusters **(Fig. 2I)**. These populations showed markedly different patterns of clonal expansion. *GZMK*^+^ IELs contained a high proportion of large TCRαβ clonotypes, whereas the *ICOS*^+^*IL10*^+^ population showed high *GZMK* expression despite being dominated by singleton and small clonotypes **(Fig. 2I)**. Thus, elevated *GZMK* expression was not restricted to highly expanded clonotypes and could occur independently of extensive clonal proliferation.

At the disease level, mean *GZMK* expression was significantly increased in both ACD and RCD1 compared with Healthy controls (p=0.0124 and p=0.0039, respectively; **Fig. 2K**). The proportion of GZMK-positive IELs showed a stronger increase and was highest in RCD1, significantly exceeding both Healthy controls (p=0.008) and ACD (p=0.044; **Fig. 2J**). RCD2 samples retained high *GZMK* expression and a substantial GZMK-positive compartment, although neither measure increased further relative to RCD1. The investigation of tissue resident markers CD103 and CD69 across IEL clusters revealed that these populations were also phenotypically heterogeneous, and notably GZMK-high clusters expressed reduced levels of both CD103/*ITGAE* and CD69 expression (**Fig. 2I; Supplementary Fig. 3A**), consistent with remodelling of the conventional CD103-high IEL phenotype.

We next examined transcriptional changes between ACD and RCD1 within the four GZMK-high populations. Differential gene expression (DGE) analysis revealed distinct but partially overlapping changes across disease states **(Supplementary Fig. 3B; Supplementary Table 3)**. Most notably, *SIK3* expression was consistently increased in RCD1 across all four populations analysed, including *GZMK*^+^ IELs, CD8^+^*GZMK*^+^ IELs, *ICOS*^+^*IL10*^+^ IELs and NK/Tγδ cells. SIK3 has recently been implicated in the regulation of T-cell functional state, with T-cell-intrinsic SIK2/3 signalling promoting a dysfunctional immune phenotype in chronically suppressive environments [38], suggesting that recurrent *SIK3* upregulation may represent a conserved component of IEL adaptation in RCD1. Relative to Healthy controls, ACD was characterised across several of these populations by increased expression of immediate-early, stress and activation-associated genes, including *DUSP4, CREM, HSPA5, LITAF* and related response genes. In contrast, comparison of ACD with RCD1 identified a different transcriptional programme involving genes associated with adaptive regulation and cell-state maintenance, including *BACH2, TOX2, STAT4, FOXO1* and *KLRK1*; intracellular signalling, including *CAMK1D, PIK3R5* and *PLCL1*; metabolic and cellular fitness, including *SIK3, RPTOR, ATP6V0C, XPR1* and *GGCT*; and cytoskeletal remodelling, including *ARHGAP15* and *RAPGEF6*. These changes were particularly concordant between CD8^+^*GZMK*^+^ and NK/Tγδ populations, while *GZMK*^+^ and *ICOS*^+^*IL10*^+^ IELs displayed partially overlapping but population-specific responses. Conversely, selected genes associated with inhibitory signalling and lineage or tissue-associated identity were reduced in RCD1 in specific populations, including *PDCD1* in CD8^+^*GZMK*^+^ and NK/Tγδ cells, *GNLY* and *TRDC* in *GZMK*^+^ and CD8^+^*GZMK*^+^ IELs, and *CXCL16* and *TMIGD2* in *ICOS*^+^*IL10*^+^ IELs (**Supplementary Fig. 3B**). Examination of these genes across all disease states further showed that several components of the RCD1-associated programme remained evident in RCD2, although the magnitude and composition of these changes differed between IEL populations (**Supplementary Fig. 3C**).

Finally, direct comparison of aberrant and other large TCRαβ clonotypes within *GZMK*^+^ and T effector IELs did not identify significant transcriptional differences after multiple-testing correction **(Supplementary Table 3)**. Together, these findings indicate that GZMK marks multiple clonally expanded and non-clonal IEL states and that its enrichment in RCD1 forms part of broader transcriptional remodelling extending beyond the previously identified mutation-bearing clones.

### A shared adaptive transcriptional programme underlies IELs in RCD1

The combination of IEL clones carrying driver mutations across multiple immune cell types, together with the marked heterogeneity observed among conventional and unconventional IEL populations, suggests that RCD1 is characterised by coordinated remodelling of the intestinal immune compartment. We therefore systematically compared the transcriptional changes within each IEL subset utilising cluster differential gene expression analysis between ACD and RCD1 (**Supplementary Table 3**).

Across the immune compartment, 15 of 17 IEL populations showed recurrent enrichment of components of a shared adaptive transcriptional programme in RCD1 relative to ACD (Fig. 3A). The two exceptions were the *IL2*^+^*CCL4^+^* population and the *GZMK*^+^ IEL population, the latter containing a prominent fraction of the molecularly defined aberrant clones unique to RCD1 (Fig. 3A). This programme comprised recurrent induction of genes associated with persistent activation and adaptive survival (*BACH2* [39]*, TOX* [40] *FOXO1* [41], [42]) and negative regulator of TCR activation *CBLB,* [43], innate-like receptor and cytokine adaptation (*KLRK1,* [44]*, FCRL6* [45]*, CD70, IL12RB2, STAT4*) [46], metabolic and proteostatic fitness (*SIK3, RPTOR, SREBF2, ATP6V0C, ATG7, AMBRA1*), MTOR associated genes (RPTOR, SREBF2, AMBRA1) [47], migration and cytoskeletal remodelling (*ARHGAP15, RAPGEF6, DOCK9, ELMO1* [48]*, SH2D3C*), and chronic adaptive activation and antigen presentation (*HLA-DQB1, HLA-DQA1, PRKCQ*). In contrast, Trm, cytotoxic and proliferating IEL subsets in ACD preferentially expressed genes associated with T-cell exhaustion and checkpoint regulation (*TIGIT, PDCD1, CTLA4, HAVCR2, LAG3*) [49, 50, 51, 52], together with markers of tissue retention and chronic activation, including *CXCR6* [53]*, TNFRSF9* and *CD38* (**Fig. 3A**). These conserved transcriptional programmes were mirrored by pathway-level analyses (GSEA), which demonstrated significant enrichment of oxidative phosphorylation, MYC targets, DNA repair, E2F targets, mTORC1 signalling, cholesterol homeostasis, IL2–STAT5 signalling, and adhesion and remodelling, confirming that RCD1 is characterised by coordinated metabolic, biosynthetic and functional adaptation across the majority of IEL cell types rather than isolated activation of individual lymphocyte subsets (**Fig. 3B**). In contrast, ACD preferentially retained NK-like cytotoxicity and exhaustion/checkpoint signatures, indicating that RCD1 involves a transition from acute cytotoxic immune responses towards a tissue-wide programme of chronic adaptive remodelling (**Fig. 3B**).

**Figure 3.**
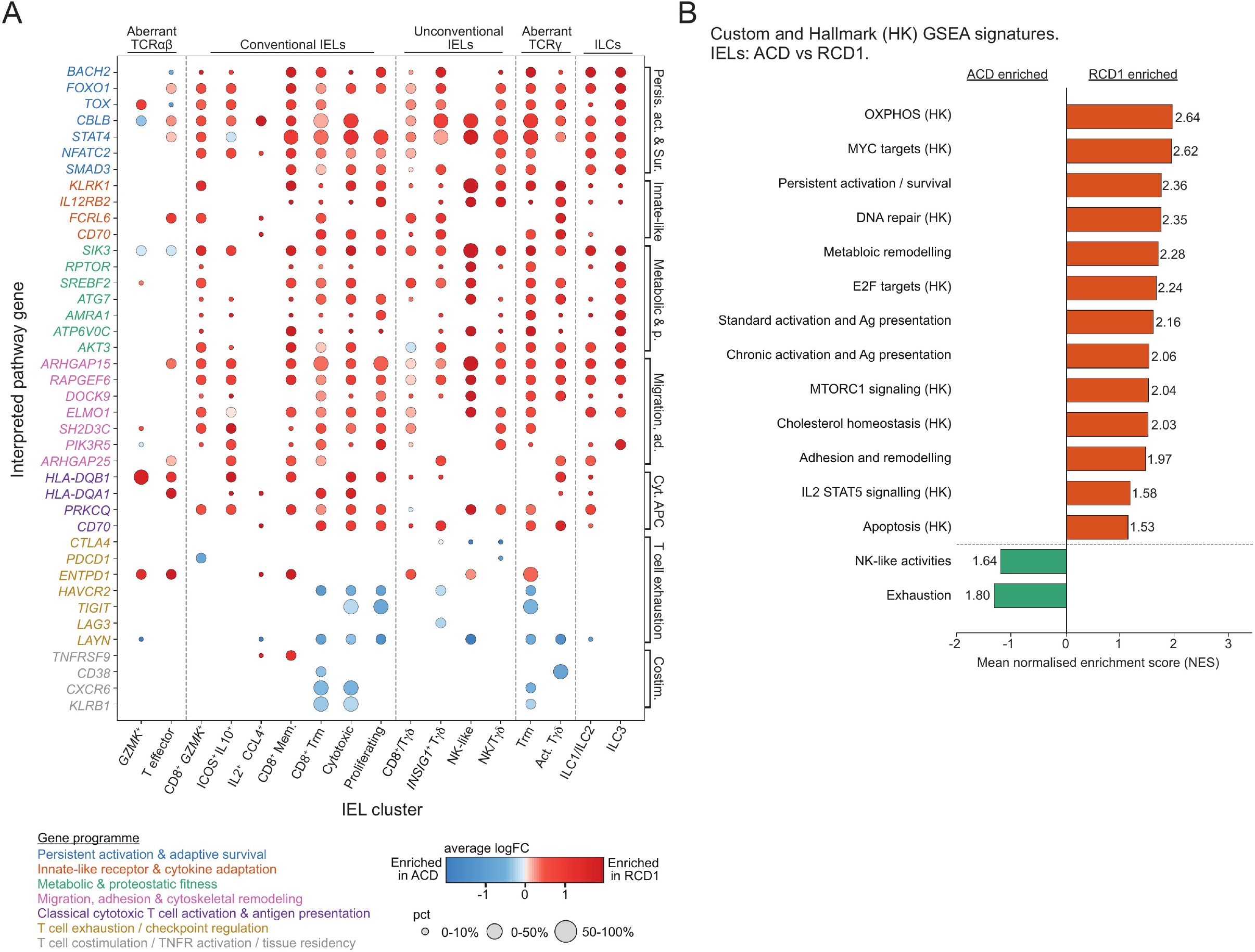
A shared adaptive transcriptional programme characterises intraepithelial lymphocytes in RCD1. **A)** Dot plot shows selected genes from within-cluster DGE analysis comparing ACD with RCD1 using edgeR across transcriptionally defined IEL populations. Genes are grouped and coloured according to functional programmes. Dot colour represents average log fold-change, with red indicating higher expression in RCD1 and blue indicating higher expression in ACD. Dot size represents the proportion of cells expressing the indicated gene within each population. **B)** Pre-ranked GSEA comparing IELs from ACD and RCD1 samples using selected Hallmark and customised gene signatures. Bars show the normalised enrichment score (NES), with positive values indicating enrichment in RCD1 and negative values indicating enrichment in ACD. ACD, active coeliac disease; GSEA, gene set enrichment analysis; IEL, intraepithelial lymphocyte; ILC, innate lymphoid cell; NES, normalised enrichment score; RCD1, refractory coeliac disease type 1; TCR, T-cell receptor.

To determine whether components of the adaptive transcriptional programme identified in RCD1 were also present in ACD, we additionally performed within-cluster differential expression between Healthy controls and ACD. ACD was characterised by recurrent enrichment of checkpoint and dysfunction-associated genes, including *CTLA4*, *PDCD1*, *ENTPD1*, *HAVCR2*, *TIGIT* and *LAYN* [49, 52] together with tissue-residency and activation-associated genes, including *CXCR6* [53], *TNFRSF9* and *CD38* (**Supp. Table 3**) [51]. In contrast, many components of the adaptive persistence, innate-like, metabolic and cytoskeletal-remodelling programme characteristic of RCD1 relative to ACD were unchanged or only sporadically altered in the Healthy versus ACD comparison. Notably, several migration-associated genes enriched in RCD1 relative to ACD, including *SH2D3C*, *RAPGEF6* and *ARHGAP25*[54, 55], were instead expressed at higher levels in Healthy than ACD IELs. These findings indicate that RCD1 is characterised by a qualitatively distinct, shared IEL programme combining adaptive survival, innate-like signalling, metabolic fitness and tissue remodelling, rather than simply amplifying the checkpoint-rich inflammatory state observed in ACD. Lineage-specific programmes are superimposed on this common adaptive state.

### Tissue remodelling characterises epithelial cells in RCD1

Given the large-scale changes in the immune compartment in refractory coeliac disease, we next investigated whether similar remodeling occurs in epithelial cells across the spectrum of coeliac disease. We performed single-cell transcriptomic profiling of EPCAM+ epithelial cells isolated from duodenal biopsies obtained from Healthy (n=6), ACD (n=5), RCD1 (n=7) and RCD2 (n=2). Unsupervised clustering identified sixteen transcriptionally distinct epithelial cell populations comprising enterocytes (*FABP1*+) transit-amplifying (TA) cells (*OLFM4*+), epithelial stem cells (*LGR5*+), enteroendocrine (EEC) (*CHGA*+), EEC progenitor cells (*TFF3*+), tuft cells (*POU2F3*+), enterocytes/colonocytes (*BEST4*+), and surface foveolar-like cells (*MUC5AC*+) (**Fig. 4A, B**). Although all major epithelial lineages were represented across disease groups, their relative abundance changed markedly in RCD (**Fig. 4C**). Among enterocyte populations, mature absorptive enterocytes predominated in healthy tissue, whereas ACD tissues were characterised by accumulation of stressed enterocytes and stressed transit-amplifying (TA) cells. RCD1 tissue was characterised by increased proportions of regenerative absorptive TA cells and enteroendocrine populations, and RCD2 by emergence of metabolically specialised epithelial populations. Tuft cells and EEC progenitors also increased from ACD to RCD1 (**Fig. 4C**). Together, these findings indicate altered epithelial differentiation and regeneration in RCD1 compared with ACD.

**Figure 4.**
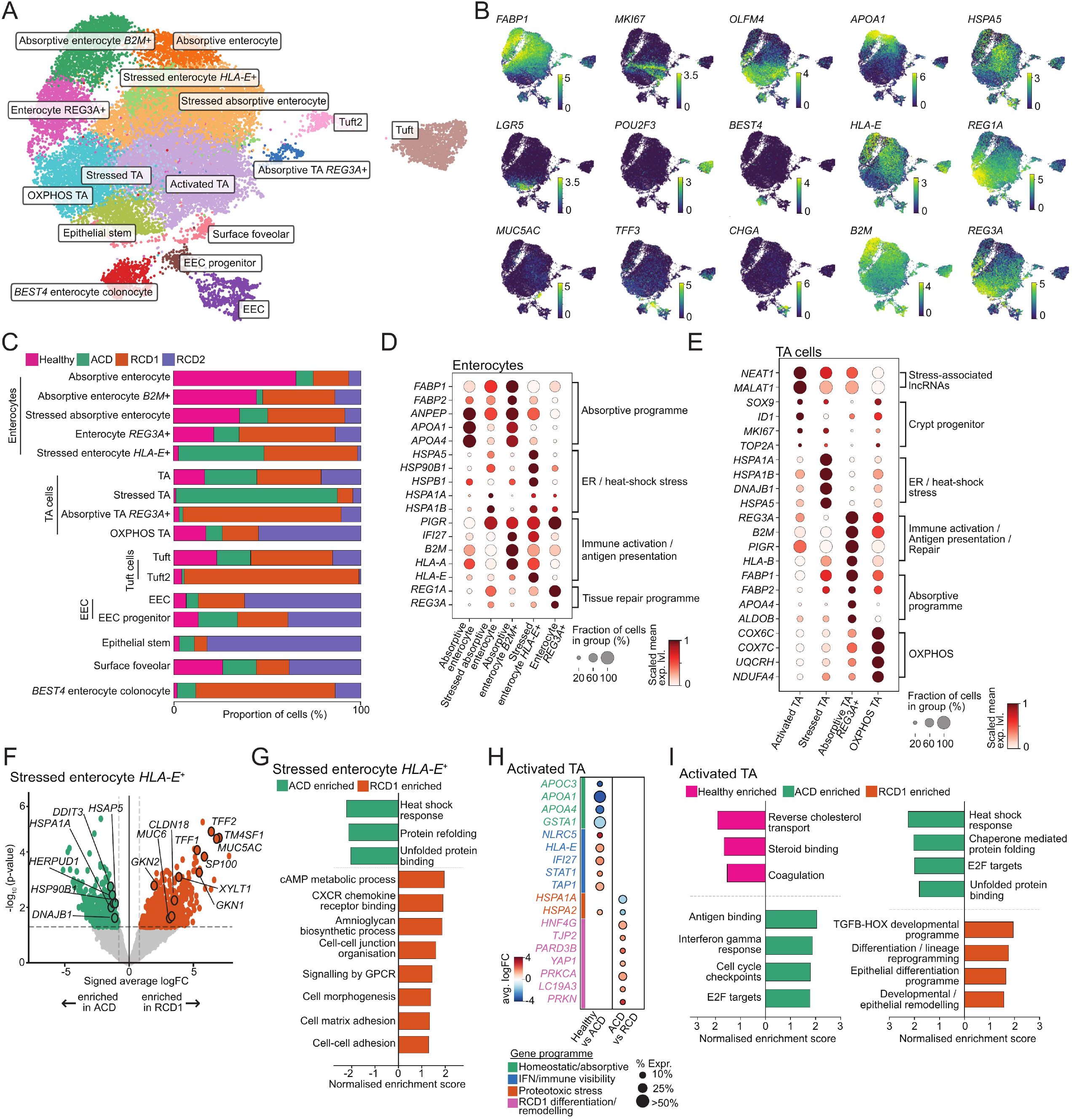
Epithelial remodelling defines distinct enterocyte and transit-amplifying cell states across coeliac disease. **A)** UMAP representation of epithelial cells from duodenal biopsies, showing 16 transcriptionally defined epithelial cell populations. **B)** UMAP representations show log-transformed normalised expression of selected marker genes used to define major epithelial lineages and disease-associated epithelial states. **C)** Stacked bar plots show the relative contribution of cells from Healthy, ACD, RCD1 and RCD2 samples to each epithelial cluster. **D-E)** Dot plots show scaled mean expression of selected genes across **D)** enterocyte and **E)** TA populations. Dot size represents the fraction of cells expressing each gene and colour represents scaled mean log-transformed normalised expression. **F)** The volcano plot shows DGE analysis between ACD and RCD1 within Stressed enterocyte *HLA-E*^+^ cluster, using pseudobulk edgeR analysis. The x-axis represents signed average log fold-change, with genes enriched in ACD shown to the left and genes enriched in RCD1 shown to the right. The y-axis represents −log_10_ *p*-value. Selected differentially expressed genes are labelled. **G)** GSEA comparing cells from ACD with RCD1 within the Stressed enterocyte *HLA-E*^+^ cluster. Bars show normalised enrichment scores (NES) for selected pathways, with negative values indicating enrichment in ACD and positive values indicating enrichment in RCD1. **H)** Dot plot showing selected differentially expressed genes from pseudobulk edgeR analyses of Activated TA cells comparing Healthy with ACD and ACD with RCD1. Genes are grouped and coloured according to functional programmes. Dot colour represents average log fold-change and dot size represents the percentage of cells expressing the indicated gene. **I)** GSEA based on within-cluster differential-expression rankings in Activated TA cells comparing Healthy with ACD (left) and ACD with RCD1 (right). Bars show NES for selected pathways, with bar direction and colour indicating the disease state in which each pathway is enriched. ACD, active coeliac disease; EEC, enteroendocrine; ER, endoplasmic reticulum; GSEA, gene set enrichment analysis; NES, normalised enrichment score; OXPHOS, oxidative phosphorylation; RCD1 and RCD2, refractory coeliac disease types 1 and 2; TA, transit-amplifying.

### Enterocytes display stress, repair and immune-interacting states across disease states

Further anlaysis of enterocytes revealed five transcriptionally distinct populations spanning mature absorptive, stress-associated, reparative and immune-interacting states (**Fig. 4A-C**). Two absorptive populations, enriched in Healthy controls, expressed canonical nutrient-absorption genes including *FABP1, FABP2, ANPEP, APOA1, APOA4, ALDOB* and *SI*, with the *B2M*^+^ population additionally expressing *HLA-A, HLA-B* and *B2M* (**Fig. 4D).** In contrast, stressed absorptive enterocytes showed prominent ER-stress and heat-shock programmes, whereas *REG3A*+ enterocytes expressed injury- and mucosal-defence-associated genes including *REG1A, REG3A, LYZ, DMBT1* and *PIGR*. Stressed *HLA-E*^+^ enterocytes displayed the strongest immune-interacting phenotype, with increased expression of classical and non-classical MHC-I molecules (*HLA-A, HLA-B, HLA-C, HLA-E*), antigen-processing genes (*B2M, TAP1, TAP2, TAPBP, PSMB8*) and interferon- and cellular-stress-associated programmes (**Fig. 4D**). Mature absorptive populations were reduced in RCD1, whereas stressed, *REG3A*+ and *HLA-E*+ states were more prominent in ACD and refractory disease (**Fig. 4C**), consistent with substantial remodelling of the absorptive epithelial compartment.

We then performed within-cluster differential expression analysis between ACD and RCD1 in Stressed enterocyte HLA-E+ cells. ACD cells preferentially expressed heat-shock and ER-stress genes, including *HSPA1A, HSPA5, HSP90B1, DDIT3, HERPUD1* and *DNAJB1*, whereas RCD1 cells showed increased gastric/foveolar-like and epithelial-plasticity genes, including *MUC5AC, TFF1, TFF2, GKN1, GKN2, MUC6, CLDN18, TM4SF1, S100P, EXT1* and *XYLT1* (**Fig 4F, Supplementary Table 4**). Consistent with these gene-level changes, GSEA showed enrichment in ACD of heat-shock, unfolded-protein-binding and protein-refolding pathways, while RCD1 was enriched for cell-junction organization, cell-cell and cell-matrix adhesion, morphogenesis, aminoglycan biosynthesis, chemokine, GPCR and CAMP signalling (**Fig 4G**).

*REG3A*+ enterocytes showed a distinct disease-associated pattern across the three clinical groups. Within-cluster edgeR analysis comparing Healthy and ACD showed significant induction in ACD of the antimicrobial and injury-response genes *REG3A, DEFA5* and *DEFA6*, with additional increases in *LYZ* and interferon-associated genes, whereas cells belonging to Healthy controls retained higher expression of absorptive and metabolic genes including *APOA1, APOC3, HSD17B2, FBP1, APOA4, FABP1* and *ALDOB* (**Supplementary Table 4**). These findings indicate that the *REG3A*+ state in ACD is characterised by loss of absorptive features and acquisition of an antimicrobial injury-response gene programme. DGE comparison between ACD and RCD1 showed reduced prominence of this antimicrobial signature in RCD1, together with increased expression of genes linked to epithelial differentiation, membrane organisation and mitochondrial function, including *HNF4G, ADH1C, EXT1, RAC1, ATP5PD, UQCRB* and *COX6C*. Consistent with these gene-level changes, GSEA of the ACD versus RCD1 comparison showed directional enrichment of antimicrobial and acute inflammatory programmes in ACD and of cytoskeletal reorganisation, wound-healing-associated cell spreading, receptor regulation and epithelial transport in RCD1, although no pathway remained significant after multiple-testing correction. Together, these analyses indicate that the *REG3A*+ population differs across disease states, with ACD displaying a prominent antimicrobial injury-response phenotype and RCD1 a more differentiated and structurally adapted programme.

Overall, enterocytes in RCD1 were distinguished by persistent stress-associated, reparative and structurally remodelled states rather than by simple amplification of the epithelial response observed in ACD. In particular, stressed *HLA-E*+ enterocytes combined epithelial remodelling with an immune-interacting phenotype that may facilitate communication with neighbouring IEL populations.

### Transit-amplifying cells diversify into disease-associated stress, regenerative and metabolic states

Transit-amplifying (TA) cells segregated into four transcriptionally distinct populations with marked differences in their distribution across disease states (**Fig. 4C**). Activated TA cells retained proliferative crypt-progenitor features, expressing *SOX9, ID1, MKI67, TOP2A* (**Fig. 4E**). In contrast, Stressed TA cells were most prominent in ACD and expressed heat-shock and unfolded-protein-response genes including *HSPA1A, HSPA1B, HSPA5, DNAJB1, DDIT3* and *HERPUD1*. The proportion of Absorptive TA REG3A+ cells were enriched in RCD1 (**Fig. 4C**) and these cells expressed high levels of the inflammation-associated regenerative genes (*REG1A, REG3A*) [56, 57], with absorptive-lineage markers (*FABP1, FABP2, APOA4*) and immune-interacting molecules including *B2M, HLA-B* and *PIGR*. OXPHOS TA cells were most prominent in RCD2 and expressed mitochondrial respiratory-chain genes including *NDUFA4, UQCRH, COX4I1, COX6C, COX7C* and *ATP5F1E*. Together, these distributions identify distinct stress, regenerative and metabolic states within the crypt compartment across disease groups.

To determine whether the Activated TA state itself changed across disease groups, we performed within-cluster differential gene expression analysis followed by GSEA comparing Healthy with ACD and ACD with RCD1 (**Fig. 4H, I, Supplementary Table 4**). Relative to Healthy controls, ACD-derived Activated TA cells showed reduced expression of genes associated with absorptive and lipid-metabolic functions, including *APOC3, APOA1, APOA4* and *GSTA1*, together with increased expression of *NLRC5, HLA-E, IFI27, STAT1* and *TAP1*, consistent with acquisition of an interferon-responsive and increasingly immune-visible phenotype. GSEA supported this transition, identifying enrichment in ACD of interferon-γ response, MHC-associated and proliferative programmes, driven by leading-edge genes including *NLRC5, IFI27, STAT1, TAP1, B2M, HLA-A* and *HLA-B*, whereas Healthy Activated TA cells showed enrichment of lipid-transport and cholesterol-associated signatures driven by *APOC3, APOA1* and *APOA4*. In the subsequent ACD versus RCD1 comparison, ACD Activated TA cells were enriched for heat-shock, protein-folding and proliferative programmes, driven by recurrent leading-edge genes including *HSPA1A, HSPA2, HSPA5, HSPH1, DNAJB1, MCM2/4/7, BIRC5* and *PLK1*, whereas RCD1-enriched signatures were driven by genes associated with epithelial differentiation and remodelling, including *HNF4G, MEIS1, HOX-family genes, SOX5, SPDEF, TGFB2, BMP5* and *TJP2*. Together, these findings indicate that Activated TA cells shift from a relatively homeostatic absorptive-metabolic state in Healthy tissue to an interferon-responsive, proliferative and proteostatically stressed state in ACD, followed by broader developmental and epithelial-remodelling programmes in RCD1.

To identify the transcriptional programme underlying the RCD1 epithelial cell subsets, we performed SCENIC analysis from both the ACD and RCD1 samples, independently (**Supp Fig 4**). SCENIC analysis revealed distinct regulatory architectures across the TA compartment. ACD-derived Activated TA cells were characterised by *HNF4A*, *ONECUT2*, *E2F3*, *E2F8* and *BRCA1* regulons associated with epithelial identity, DNA replication and cell-cycle, whereas RCD1 Activated TA cells displayed *KMT2B*, *HMG20B*, *E2F4*, *CEBPZ* and *CREBZF* regulons linked to chromatin regulation, RNA processing and mitochondrial biosynthesis. Absorptive TA REG3A+ cells were characterised by CDX1- and NR1D1-associated intestinal-lineage programmes in ACD, compared with *PHF20*, *MITF*, *BACH2* and *ZNF81* regulons in RCD1, whose targets supported cytoskeletal organisation, membrane trafficking and metabolic adaptation.

Overall, ACD TA cells were dominated by stress and proliferative programmes, whereas RCD1 showed greater regenerative and differentiation-associated states, indicating broad remodelling of the regenerative crypt compartment across disease states.

### Enteroendocrine cells acquire neurosecretory and sensory programmes in RCD1

Beyond the absorptive and progenitor compartments, enteroendocrine cells (EECs) showed marked heterogeneity across disease states (**Fig. 5A,B**). Previous single-cell studies have defined diverse hormone-producing EEC lineages, including somatostatin-producing D cells, and identified expansion of D cells and NEUROG3+ endocrine progenitors in coeliac disease[24, 58]. Guided by these markers, sub-clustering resolved two major states: a sensory-secretory (S) population, expressing *CCK, GCG* and *NTS* together with sensory-associated genes, and a somatostatin-producing D-cell population defined by high *SST* expression (**Fig. 5A,C**). Sample-level abundance analysis showed increased representation of the EEC compartment in refractory disease, with both populations more abundant in RCD1 than in Healthy and ACD samples (**Fig. 5B**). The increase was particularly pronounced for D cells, which were sparse in Healthy and ACD tissues but enriched in refractory samples.

**Figure 5.**
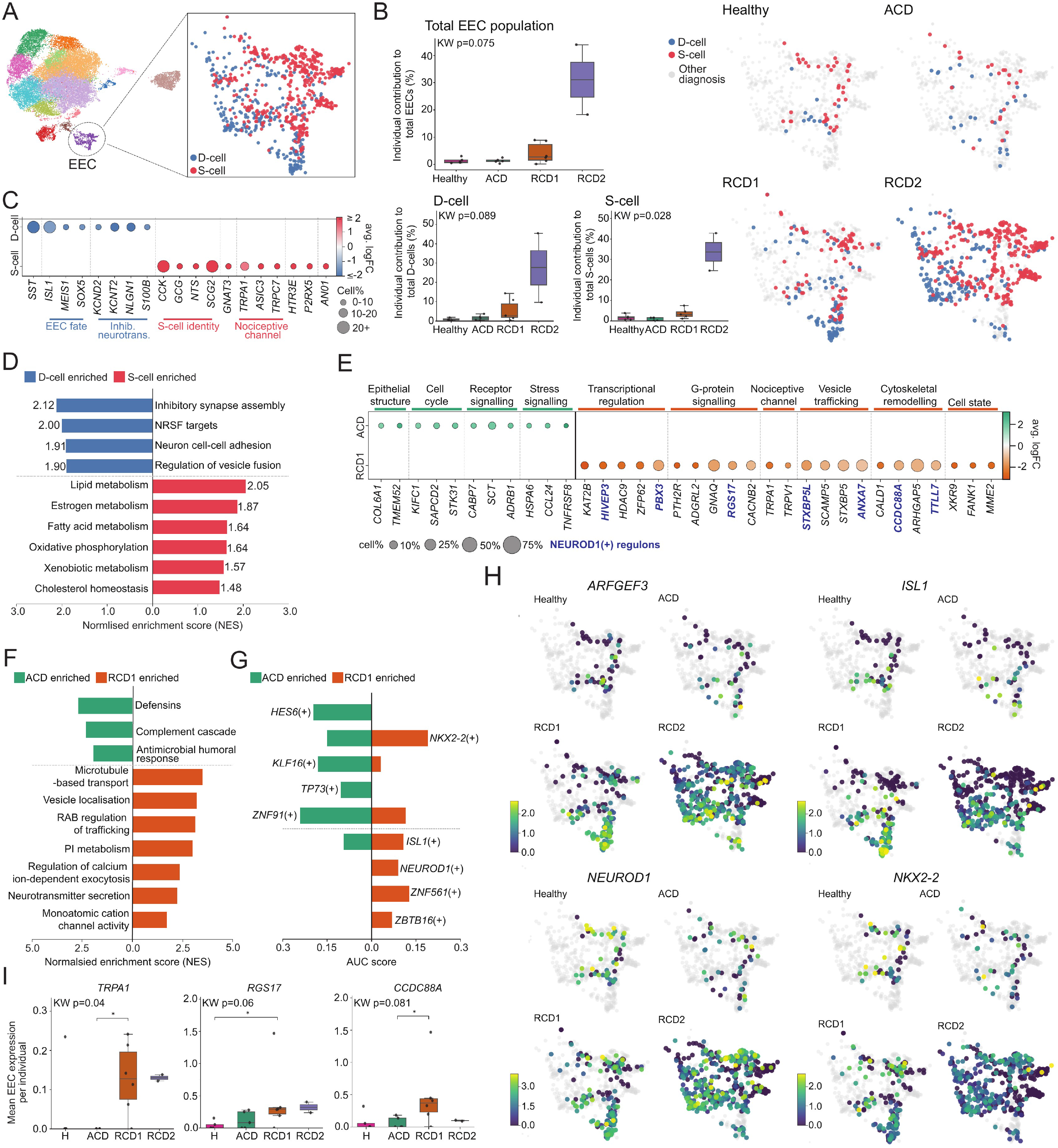
Enteroendocrine cells expand and acquire specialised sensory and neurosecretory programmes in RCD. **A)** UMAP representation of epithelial cells highlighting the enteroendocrine cell (EEC) population, subclustering them into D-cell and S-cell states. **B)** Boxplots show the contribution of total EECs, D-cells and S-cells to the epithelial compartment across Healthy, ACD, RCD1 and RCD2 samples. Each point represents an individual sample. Kruskal–Wallis (KW) *p*-values are shown. UMAPs show the distribution of D- and S-cells within the EEC compartment for each disease state. **C)** Dot plot shows selected genes distinguishing D- and S-cell states. Dot size represents the percentage of cells expressing each gene and colour represents scaled mean log-transformed normalised expression. **D)** GSEA comparing D-cells and S-cells within EEC population. Bars represent normalised enrichment scores (NES). **E)** The dot plot shows selected differentially expressed genes in EECs between ACD and RCD1 samples using edgeR. Dot size represents the percentage of cells expressing each gene and colour represents average log fold-change. Genes associated with the NEUROD1 regulon are indicated in blue. **F)** GSEA of DGE between EEC cells from ACD and RCD1 samples showing selected pathways enriched in ACD or RCD1. Bars represent NES. **G)** SCENIC gene regulatory network analysis showing selected regulons differentially enriched in EECs from ACD or RCD1 samples. Bars represent AUC scores. **H)** UMAP representations show the expression of selected transcriptional regulators identified through SCENIC analysis across EECs from Healthy, ACD, RCD1 and RCD2 samples. Colour represents mean log-transformed normalised gene expression. **I)** Boxplots show sample-level mean expression of selected genes in EECs across Healthy, ACD, RCD1 and RCD2 samples. Each point represents an individual sample. Kruskal–Wallis *p*-values are shown, with the indicated significant pairwise comparisons shown where applicable. ACD: active coeliac disease; EEC: enteroendocrine cell; GSEA: gene set enrichment analysis; KW: Kruskal–Wallis; NES: normalised enrichment score; AUC: area under curve; RCD1 and RCD2: refractory coeliac disease types 1 and 2.

To define the functional differences between these populations, we compared S and D cells by differential expression and GSEA (**Fig. 5C,D**). S cells preferentially expressed enteroendocrine hormones (*CCK, GCG, NTS*) together with sensory and nociceptive genes (*GNAT3, TRPA1, ASIC3, TRPC7, HTR3E, P2RX5, ANO1*). In contrast, D cells expressed *SST* and lineage-associated transcription factors (*ISL1, MEIS1, SOX5*), together with genes involved in membrane excitability and neuronal communication (*KCND2, KCNT2, NLGN1, S100B*). GSEA further distinguished these states, with D cells enriched for inhibitory synapse assembly, neuronal cell–cell adhesion and vesicle-fusion pathways, whereas S cells showed enrichment of lipid, fatty-acid and oxidative metabolic programmes. These analyses identify a sensory-secretory S-cell state and a distinct neuromodulatory D-cell programme.

We next examined disease-associated transcriptional differences across the broader EEC lineage by analysing EEC progenitors and differentiated EECs together using edgeR. Compared with Healthy controls, ACD showed increased expression of *SPINK4, SHISAL2B* and *SCGB2A1*, together with progenitor, secretory-lineage and stress-associated genes including *MKI67, SPDEF, TFF2, AGR2, HSP90B1, IFI27* and *AREG*, whereas *PYY* was higher in Healthy cells (**Supplementary Table 4**). Thus, ACD was characterised by a regenerative, secretory and stress-associated EEC programme distinct from the more mature endocrine profile observed in Healthy tissue.

Comparison of ACD and RCD1 using DGE (edgeR) and GSEA revealed a distinct RCD1-associated programme (**Fig. 5E, F**). RCD1 showed coordinated increases in genes associated with nociceptive and sensory signalling, including *TRPA1, TRPV1, CACNB2, GNAQ* and *RGS17*. This was accompanied by increased expression of genes involved in regulated secretion and vesicle trafficking (*STXBP5, STXBP5L, SCAMP5*) and cytoskeletal and membrane organisation (*CCDC88A, TTLL7, PBX3, ARHGAP5*). Consistent with these gene-level differences, GSEA showed enrichment in RCD1 of microtubule-based transport, vesicle localisation, RAB-regulated trafficking and PI metabolism, together with calcium-ion dependent exocytosis, neurotransmitter secretion and ion-channel activity. In contrast, ACD was enriched for defensin, complement and antimicrobial-response programmes. Together, these findings identify an RCD1-associated nociceptive-neurosecretory programme, integrating enhanced sensory signalling with intracellular trafficking and regulated secretion.

We next used SCENIC to define regulatory programmes associated with EEC remodelling in ACD and RCD1 (**Fig. 5G, H**). ACD EECs were characterised by regulons including *HES6*, *KLF16*, *TP73* and ZNF91, whereas RCD1 showed prominent activity of *NKX2-2*, *ISL1* and, notably, *NEUROD1* (**Fig. 5G**). Mapping regulon activity across EECs further demonstrated greater *NEUROD1* and *ISL1* activity in refractory samples compared with Healthy and ACD tissues (**Fig. 5H**). Consistent with the edgeR and GSEA results, RCD1 samples also showed increased expression of sensory and signalling-associated genes including *RGS17*, *CCDC88A* and the nociceptive channel gene *TRPA1* (**Fig. 5I**). Together, these analyses support an RCD1-associated EEC state characterised by NEUROD1-centred regulatory activity, enhanced sensory signalling and neurosecretory remodelling.

Overall, RCD1 was characterised by expansion and transcriptional remodelling of specialised EEC states, together with a NEUROD1-centred nociceptive-neurosecretory programme. These findings implicate the enteroendocrine compartment in altered epithelial-neural and neuroimmune communication during chronic mucosal inflammation.

### Spatial transcriptomics localises epithelial remodelling and immune–epithelial interactions in refractory coeliac disease

To determine the anatomical distribution of the epithelial and IEL states identified by scRNA-seq, we performed Visium spatial transcriptomics across six duodenal biopsies from Healthy (n=1), ACD (n=1; two tissue sections), RCD1 (n=2) and RCD2 (n=1) individuals (**Fig. 6; Supplementary Fig. S5**). Spatial gene expression was integrated with histological annotation to define regions of interest (ROIs) representing distinct epithelial and immune microenvironments.

**Figure 6.**
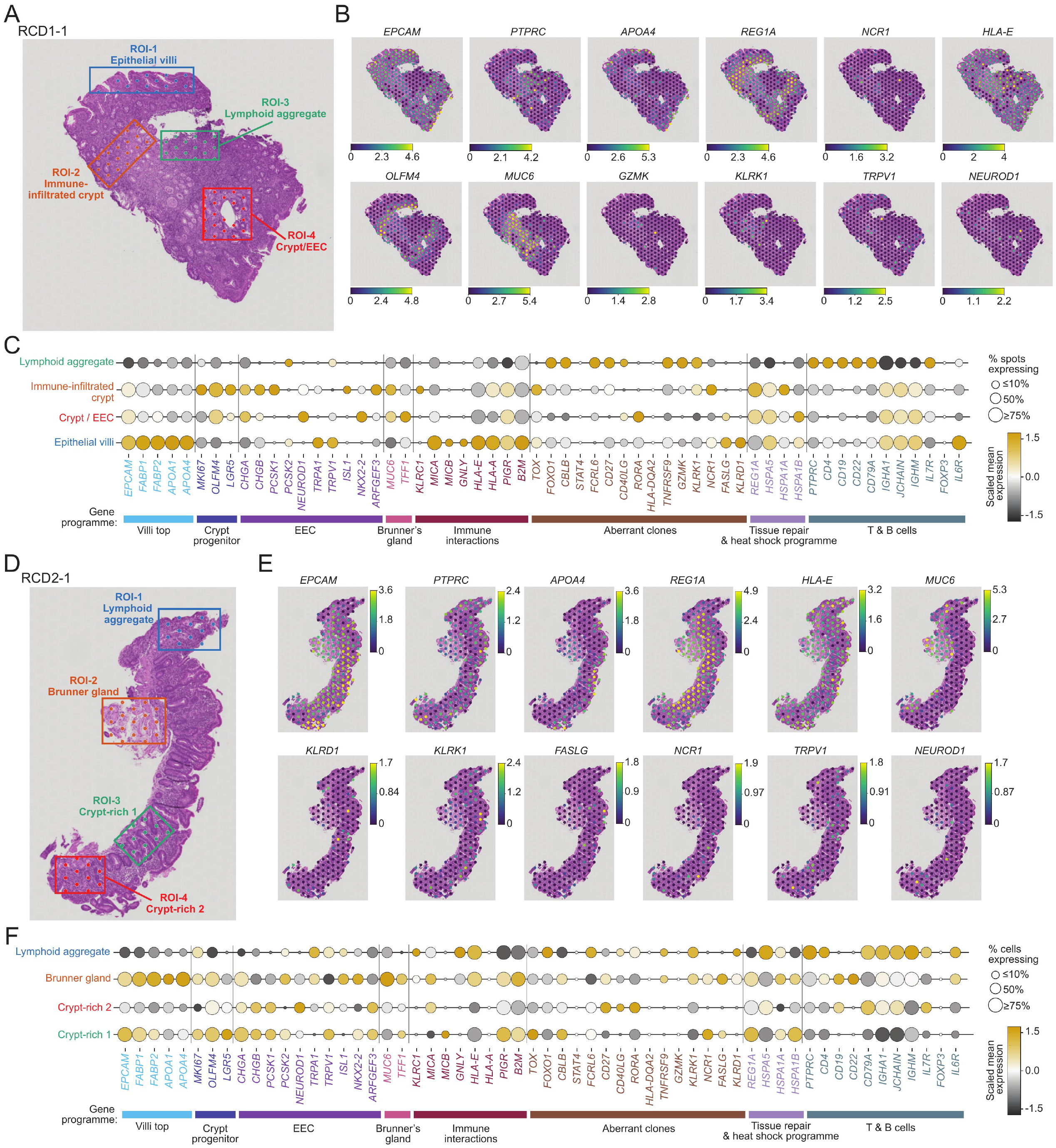
Spatial transcriptomics identifies localised epithelial–immune microenvironments in RCD. **A)** H&E-stained tissue section from an RCD1 (RCD1-1) biopsy showing four histology- and marker-guided regions of interest (ROIs) **B)** Spatial expression maps of selected genes across the RCD1 tissue section. Colour represents log-transformed normalised gene expression at individual Visium spots. **C)** The dot plot shows selected ROI-associated genes. For each ROI, gene expression was compared with the remaining ROIs to identify transcriptional features enriched within each spatial microenvironment. Selected genes are grouped and coloured according to programmes. Dot size represents the percentage of Visium spots within each ROI expressing the indicated gene, and colour represents scaled mean log-transformed normalised expression across spots within each ROI. **D)** H&E-stained tissue section from an RCD2 (RCD2-1) biopsy showing four histology- and marker-guided ROIs. **E)** Spatial expression maps of selected genes across the RCD2 tissue section. Colour represents log-transformed normalised gene expression at individual Visium spots. **F)** The dot plot shows selected ROI-associated genes. For each ROI, gene expression was compared with the remaining ROIs to identify transcriptional features enriched within each spatial microenvironment. Genes are grouped and coloured according to the same biological programmes shown in **C**. Dot size represents the percentage of Visium spots within each ROI expressing the indicated gene, and colour represents scaled mean log-transformed normalised expression across spots within each ROI. ACD, active coeliac disease; EEC, enteroendocrine; H&E, haematoxylin and eosin; IEL, intraepithelial lymphocyte; RCD1 and RCD2, refractory coeliac disease types 1 and 2; ROI, region of interest.

Histological annotation of the RCD1-1 biopsy identified four spatially distinct microenvironments comprising villus/surface epithelium, a crypt/EEC-enriched region, an immune-infiltrated crypt and a lymphoid aggregate (**Fig. 6A**). Spatial expression maps showed the anatomical distribution of representative epithelial, regenerative, EEC and immune-associated genes across the tissue section (**Fig. 6B**). Differential gene expression analysis between the 4 ROIs, revealed distinct transcriptional profiles (**Fig. 6C**). The villus ROI was dominated by mature absorptive markers, with high and broadly distributed expression of *EPCAM, FABP1, APOA4* and *MKI67*, together with prominent *HLA-E* expression, whereas the crypt/EEC region showed reduced absorptive identity and enrichment of crypt-progenitor and enteroendocrine programmes, including *OLFM4, NEUROD1* and *PCSK1/2* (**Fig. 6C**). Notably, the immune-infiltrated crypt retained these epithelial/EEC features while also expressing immune-visibility and innate-like lymphocyte-associated genes, including *HLA-E, MICA* and *KLRK1*. In contrast, the neighbouring lymphoid aggregate was dominated by lymphocyte-associated genes and also expressed *GZMK*, *NCR1*, and *KLRK1* expression, consistent with the presence of activated/innate-like lymphocyte populations, which were shown to be over expressed in the aberrant clone identified in this individual (Fig. 1) [18]. A second biopsy (RCD1-16) showed a similar focal microenvironment in which regenerative and EEC-associated epithelial programmes spatially overlapped with infiltrating T-cell and innate-like IEL signals, including *KLRK1* and *ITGAE* **(Supplementary Fig. 5A-C)**. Together, these findings identify focal RCD1 mucosal regions in which regenerative and immune-infiltrated microenvironment are spatially juxtaposed with innate-like IEL populations.

A similar spatial organisation was observed in RCD2 biopsy (**Fig. 6D-F**). Two crypt-rich regions, a Brunner’s gland region (expressing *MUC6*) and a lymphoid aggregate showed distinct combinations of epithelial, regenerative and immune programmes (**Fig. 6D, E**). Crypt-rich regions retained progenitor/EEC-associated features together with *REG1A*-associated tissue-remodelling signatures.The Brunner’s gland ROI showed prominent epithelial and absorptive profiles, including *EPCAM, FABP1, FABP2, APOA1* and *APOA4*. This region also expressed crypt/EEC-associated genes (e.g. *CHGA, NEUROD1, TRPA1, NKX2-2*), and regenerative and immune-visibility associated genes such as *REG1A, HLA-E, HLA-A, B2M* and *PIGR*. This region also contained detectable innate-like lymphocyte signals (**Fig. 6F**). In contrast, the lymphoid aggregate was broadly enriched for expression of T- and B-cell-associated genes (*CD3D, CD4, CD19, CD22, IL7R* and *FOXP3)*. This region also showed prominent *KLRK1* and *GZMK*, together with *HLA-E*, consistent with an activated lymphoid compartment containing innate-like lymphocyte populations. Epithelial and absorptive markers were markedly reduced relative to the Brunner’s gland region, although *TRPA1* and some EEC-associated signals remained detectable. These data therefore identified discrete RCD2 regions in which regenerative and immune-visible epithelial states were spatially juxtaposed with innate-like lymphocyte populations, consistent with spatial proximity between altered epithelial compartments and infiltrating innate-like IELs.

These focussed epithelial–immune microenvironments observed in RCD1 and RCD2 were less evident in the samples Healthy and ACD tissues (**Supplementary Fig. S5D-G**), consistent with previous spatial profiling of coeliac disease [24]. Healthy tissue retained relatively organised absorptive, crypt/progenitor and EEC-associated compartments (**Supplementary Fig. S5D, E**). In contrast, ACD showed substantial epithelial remodelling, including prominent *REG1A* expression, altered absorptive differentiation and *CHGA*- and *MUC6*-positive epithelial regions (**Supplementary Fig. S5F, G)**. Although immune-cell infiltration was detectable in ACD, the focal juxtaposition of regenerative and EEC-associated epithelium with *KLRK1*-expressing innate-like IEL signals was less apparent than in RCD.

Overall, these spatial transcriptomic data supported the anatomical localization of the epithelial and IEL programmes identified scRNA-seq within discrete mucosal microenvironments. In RCD1 and RCD2, regenerative and immune-visible epithelial programmes were spatially juxtaposed with KLRK1/GZMK-expressing innate-like IEL populations, whereas Healthy and ACD tissues showed less pronounced epithelial-immune convergence. These findings provide spatial evidence that refractory disease is associated with focal epithelial-immune microenvironment that may sustain chronic mucosal pathology.

## Discussion

Our study identifies coordinated remodelling of both the immune and epithelial compartments in RCD1 and defined molecular features that distinguish refractory disease from the inflammatory response established in active coeliac disease. RCD1 was characterised by widespread clonal expansion across conventional and unconventional αβ and γδ IEL populations, extending beyond previously identified mutation-bearing aberrant clones, together with a shared transcriptional programme integrating adaptive persistence, innate-like signalling, metabolic fitness and tissue remodelling. In parallel, the epithelial compartment underwent extensive reorganisation, including stress-associated and reparative enterocyte states, altered regenerative programmes within the crypt compartment, and expansion of specialised enteroendocrine populations with sensory and neurosecretory features. Spatial transcriptomics further localised these epithelial and immune programmes within focal mucosal microenvironments in refractory tissue. Together, these findings define RCD1 as a distinct mucosal state in which broad IEL clonal and transcriptional adaptation is accompanied by regenerative, immune-interacting and neurosecretory epithelial remodelling, rather than simply representing as amplification of ACD. This model does not imply a linear progression from RCD1 to RCD2 or EATL, but instead identifies molecular processes associated with refractory disease that may contribute to its heterogeneous clinical trajectories.

A major finding was that RCD1 involved coordinated remodelling of both IEL transcriptional state and TCR clonality. Across multiple conventional and unconventional IEL populations, RCD1 was characterised by a shared programme of chronic activation and persistence, innate-like receptor and cytokine signalling, and metabolic and proteostatic adaptation, including *KLRK1, IL12RB2, FCRL6, RPTOR, SREBF2, ATG7* and *AMBRA1*. This differed from the checkpoint-, exhaustion- and tissue-residency-associated programmes more prominent in ACD and extended beyond the somatically mutated aberrant clones. In parallel, expanded TCRαβ and TCRγδ clonotypes were distributed across multiple IEL populations rather than being confined to molecularly defined aberrant clones, with individual expanded clonotypes occupying distinct transcriptional states. This suggests that clonal selection represents a broader feature of the refractory immune compartment, upon which somatic mutations and lineage-specific programmes may subsequently impose additional functional diversity. The localisation of individual large clonotypes across related transcriptional states further supports considerable phenotypic plasticity within clonally expanded IEL populations.

However, the cross-sectional design does not establish the temporal relationship between clonal expansion, somatic evolution and transcriptional remodelling. GZMK IELs represented a prominent component of the remodelled immune landscape in RCD1, spaning both clonally expanded and non-expanded populations. These cells combined effector-associated features with increased CD69, and comparatively reduced CD103 across several IEL subsets, indicating that the GZMK phenotype is not restricted to mutation-bearing or highly expanded clones. GZMK T-cell states have also been described in other settings of chronic immune activation [35, 36, 37], while recent single-cell studies of RCD2 and EATL identified GZMK-expressing tissue-resident T-cell populations [26]. Although the relationship between these states across chronic coeliac inflammation and lymphoma remains unresolved, their recurrence supports further investigation of GZMK IELs as a feature of persistent mucosal immune remodelling.

Our data indicate that the epithelial compartment undergoes extensive remodelling in RCD, extending changes previously described in ACD, including loss of mature absorptive programmes and emergence of inflammatory and regenerative epithelial states [24]. This was particularly evident in the Stressed enterocyte HLA-E+ population, which combined MHC class I and interferon-response programmes with broader structural and signalling remodelling. HLA-E therefore appears to mark an epithelial state with increased immune visibility rather than representing an isolated disease-associated gene. This is relevant because intestinal epithelial cells can engage cytotoxic IELs through both MHC- and stress-dependent pathways. Recent work in ACD demonstrated IFNγ-enhanced, TCR-dependent recognition and killing of villous enterocytes expressing HLA-E and HLA-B [59], while epithelial MICA/B can engage NKG2D/KLRK1 on IELs during epithelial stress [19, 21]. Together, these observations suggest that epithelial remodelling in RCD1 may alter the interface between stressed epithelium and increasingly innate-adapted IEL populations. Spatial transcriptomics provided complementary support for this model by localising epithelial and immune programmes within discrete mucosal microenvironments. In RCD1 and RCD2, regenerative and immune-visible epithelial programmes, including *REG1A* and *HLA-E*, occurred in proximity to lymphoid-rich regions expressing *KLRK1* and *GZMK*-associated innate-like lymphocyte signals. These patterns were less evident in the sampled Healthy and ACD tissues and were broadly consistent with previous spatial studies of coeliac disease [24]. Although limited by sample size and the multicellular resolution of Visium, these findings support the presence of focal epithelial–immune microenvironments in refractory disease.

Our study extends epithelial remodelling beyond mature enterocytes to progenitor and secretory compartments. Whereas ACD TA cells were dominated by proliferative and stress-associated programmes, RCD1 TA states showed greater regenerative and differentiation-associated features, consistent with altered epithelial repair. More unexpectedly, RCD1 EECs acquired a coordinated sensory and neurosecretory programme involving *TRPA1, TRPV1*, nociceptive ion-channel and calcium signalling, regulated secretion and prominent NEUROD1-associated regulatory activity. EECs can act as epithelial sensory transducers and communicate with peripheral neurons through neuroactive mediators and sensory pathways, including TRPA1 [60, 61]. Experimental manipulation of enterochromaffin-cell activity has further shown that sustained activation can promote visceral hypersensitivity and pain-related responses [62]. These findings raise the possibility that EEC remodelling in RCD affects not only epithelial barrier and immune functions, but also epithelial–neural and neuroimmune communication. Whether these altered EEC states contribute directly to persistent symptoms or mucosal inflammation will require functional investigation.

Several limitations should be considered. Most importantly, the cross-sectional design precludes determination of the temporal sequence linking ACD and RCD1, and longitudinal sampling will be required to establish whether the immune and epithelial states identified here precede refractory disease or reflect its established tissue environment. The small RCD2 cohort also limits assessment of heterogeneity within this subtype. Recent work by Malamut and colleagues provides complementary evidence that clonal evolution and JAK-STAT dysregulation are shared features across RCD1 and RCD2 [26]. They identified large clonal cytotoxic CD8+ T-cell expansions in RCD1 carrying somatic alterations affecting the JAK-STAT pathway, including loss-of-function mutations in SOCS1 and SOCS3, independently supporting our previous identification of somatically mutated aberrant clones in RCD1 [18], while also demonstrating substantial genomic and transcriptional heterogeneity within RCD2 tumour populations.

Our study extends recent observations of clonal lymphocyte evolution in refractory coeliac disease by defining the broader immune and epithelial landscape associated with RCD1. Increased TCR clonal expansion across multiple αβ and γδ IEL states was accompanied by shared adaptive transcriptional programmes and extensive remodelling of progenitor, absorptive and specialised epithelial compartmsents. Together, these findings define RCD1 as a state of coordinated immune and epithelial remodelling in which clonal lymphocyte expansion occurs within a broader mucosal ecosystem characterised by IEL adaptation and regenerative, immune-visible and neurosecretory epithelial programmes. Rather than representing simply intensified ACD, RCD1 therefore appears to involve a qualitatively distinct mucosal state in which persistent immune activation and epithelial adaptation converge and may contribute to chronic tissue injury.

## Materials and Methods

### Study design

The objective of this study was to define immune and epithelial cell states associated with refractory coeliac disease, with a particular focus on distinguishing RCD1 from ACD. Duodenal biopsies were obtained from individuals with RCD1, RCD2, newly diagnosed ACD and healthy controls. Immune (CD45) and epithelial (EPCAM) compartments were isolated in parallel and profiled using CITE-seq and scRNA-seq respectively. Immune cells were additionally profiled using TCR sequencing, enabling integration of transcriptional, phenotypic and clonal information at single-cell resolution.

Sample size was primarily determined by the availability of clinically well-characterised human biopsy material, particularly for RCD. The study was observational and cross-sectional and therefore randomisation was not applicable. Single-cell data were subjected to predefined quality control procedures, and all biological samples passing sample-level quality control criteria were retained. Where possible, major findings were evaluated using complementary approaches including differential gene expression, pathway enrichment, TCR repertoire, transcription factor regulon and spatial transcriptomic analyses.

### Patient enrolment and sample collection

Diagnosis of coeliac disease was based on established clinical criteria [4], including positive coeliac serology, together with compatible duodenal histopathology. RCD was defined by persistent or recurrent malabsorptive symptoms and villous atrophy despite more than 12 months of a strict gluten-free diet, with dietary adherence assessed by expert dietetic review and alternative causes of enteropathy excluded by clinical investigation. RCD2 was diagnosed by demonstration of an aberrant IEL population >20% of the total IELs [8, 63], using immunohistochemistry and flow cytometry. This included detection of an expanded aberrant CD45 CD103 sCD3^−^ cytoplasmic-CD3^+^ CD8^−^ IEL population by flow cytometry according to established diagnostic thresholds [18]. RCD1 was diagnosed in individuals fulfilling clinical and histological criteria for refractory disease but lacking the aberrant IEL phenotype diagnostic of RCD2 and after exclusion of alternative causes of enteropathy. ACD samples were obtained from individuals with positive coeliac serology undergoing diagnostic endoscopy, with subsequent histopathological confirmation of coeliac disease, before commencement of a gluten-free diet. Healthy controls were individuals undergoing endoscopy in whom ACD was excluded by negative coeliac serology and normal or near-normal duodenal histology (Marsh score <1). Clinical and demographic characteristics are provided in **Supplementary Table 1**.

Duodenal biopsies were obtained during routine upper gastrointestinal endoscopy at the Royal Melbourne Hospital and Melbourne Private Hospital (Melbourne, Australia), Blacktown Hospital (Sydney, Australia), and Fondazione IRCCS Policlinico San Matteo (Pavia, Italy). Typically, four to six biopsies were obtained from the second part of the duodenum and two from the duodenal bulb. The biopsies from the second part of the duodenum were placed immediately in ice-cold RPMI-1640, washed twice in phosphate-buffered saline (PBS), transferred to CryoStor CS10 freezing medium (STEMCELL Technologies) and stored in liquid nitrogen. Remaining biopsies were processed for routine histopathological assessment and Marsh grading [4].

### Duodenal biopsy dissociation

Cryopreserved biopsies were thawed in a 37°C water bath and washed twice with RPMI-1640. each tissue was incubated twice with 2 mM EDTA for 15 min at 37°C with continuous rotation. Supernatants containing epithelial fractions were combined, washed and maintained on ice.

The remaining tissue was digested with 1 U/mL Collagenase IV (STEMCELL Technologies, Cat. 7426) and 0.05 mg/mL DNase I (Sigma, Cat. 10104159001) in RPMI-1640 for 1 h at 37°C with continuous rotation. Following digestion, residual tissue was mechanically dissociated by repeated passage through an 18-gauge needle and combined with the epithelial fraction. The resulting suspension was filtered through a 100-µm cell strainer, and cell number and viability were determined using Trypan Blue exclusion.

### Flow cytometric isolation of immune and epithelial cells

Single-cell suspensions were resuspended in PBS containing 2% fetal calf serum (FCS) and incubated with Human TruStain FcX (BioLegend) for 15 min at 4°C. Cells were subsequently stained for 30 min at 4°C with PE-conjugated anti-human CD45 (BioLegend, clone HI30, Cat. 304008) and FITC-conjugated anti-human EPCAM/CD326 (BioLegend, clone 9C4, Cat. 324203). Antibodies were used at a 1:50 dilution. Cells were washed twice in PBS containing 2% FCS. CD45 immune cells and EPCAM epithelial cells were sorted separately in bulk by fluorescence-activated cell sorting on a FACS Aria III (BD Biosciences). Dead cells were excluded using DAPI staining.

### Single-cell RNA sequencing and CITE-seq

Flow-sorted CD45 cells were blocked with Human TruStain FcX for 15 min on ice and subsequently incubated with a panel of 159 TotalSeq-A oligonucleotide-conjugated antibodies (BioLegend) for 30 min on ice. The complete custom-made antibody panel is provided in **Supplementary Table 2**. Following three washes in PBS containing 2% FCS, cells were filtered through a 40-µm strainer and counted using a haemocytometer.

Single-cell RNA libraries were generated using the Chromium Single Cell 3′ v3 platform (10x Genomics) according to the manufacturer’s protocol, targeting recovery of approximately 10,000–15,000 cells per sample. CD45 and EPCAM cells were processed separately. For immune samples, oligonucleotide-tagged antibody libraries were generated in parallel with gene expression libraries.

Libraries were sequenced using an Illumina NovaSeq instrument with 150-bp paired-end reads, targeting approximately 50,000 reads per cell for gene expression libraries and approximately 30,000 reads per cell for antibody-derived tag libraries. Full-length TCRα, TCRβ, TCRγ and TCRδ sequences were obtained from the single-cell RNA libraries using RAGE-seq [29]. Between 50-100 ng of full-length cDNA was used for targeted enrichment of all functional TCR and BCR genes with the xGen NGS Hybridization Capture protocol (Integrated DNA Technologies), according to the manufacturer’s recommendations.

### Single-cell gene expression and surface protein data preprocessing

Gene expression matrices containing unique molecular identifier (UMI) counts were generated using Cell Ranger v3.0.1 (10x Genomics), with alignment to the GRCh38 human reference genome. CITE-seq antibody-derived tag count matrices were generated using CITE-seq-Count [64]. Hashtag oligonucleotide (HTO) count matrices were CLR-normalised and used for sample demultiplexing with Seurat’s *HTODemux()* function. Cells were subjected to quality control filtering based on mitochondrial transcript percentage, number of detected genes and total UMI counts. Manual thresholds were applied separately to each sample and cells failing any criterion were excluded. Following QC, RNA counts were normalised using scran [65] and further log1p transformed where applicable and protein counts were normalised using centred-log-ratio transformation implemented in Seurat.

### Integration, dimensionality reduction and clustering

The normalised gene expression values were then subjected to batch correction. For immune and epithelial cells separately, 4000 most highly variable genes were identified using Seurat_v3 method implemented in Scanpy [66], with sample identity specified as the batch variable. The scVI integration method was used to mitigate the batch effects [67], with the model being configured with 2 hidden layers, a 30-dimensional latent space and a negative binomial gene expression likelihood. The model was trained for a maximum of 40 epochs with early stopping, and used to construct the nearest neighbour graph and generate UMAP embeddings with the UMAP minimum distance set to 0.3. Leiden clustering [68] was then performed at multiple resolutions (ranging from 0-5), with the 1.5 and 0.8 values being selected based on cluster transcriptional for corresponding immune and epithelial cells, and additional surface protein markers, and where relevant TCR information for immune cells, respectively.

### TCR reconstruction using RAGE-seq and clonotype analysis

T-cell receptor sequences were reconstructed using the RAGE-seq framework as previously described [29]. RAGE-seq enables recovery of antigen-receptor sequences while preserving linkage to the corresponding single-cell transcriptomic barcode, allowing integration of TCR sequence with transcriptional and phenotypic information.

Productive TCRαβ and TCRγδ sequences were assigned to individual cells using single-cell barcodes. TCR chains were annotated for V, D and J gene usage and complementarity-determining region 3 (CDR3) sequence. Cells containing productive paired TCRαβ or TCRγδ sequences were retained for repertoire analyses. Clonotypes were defined as cells containing unique paired CDR3 amino acid sequences.

Clone size was calculated as the number of cells sharing a clonotype within each sample. Repertoire diversity was quantified using normalised Shannon entropy, calculated as the Shannon entropy divided by the logarithm of clonotype richness. Clonal sharing between cell populations was quantified using the Jaccard index, defined as the number of shared clonotypes divided by the total number of unique clonotypes across both populations. All TCR analyses were performed in R using custom functions.

### Differential gene expression analysis

Differential gene expression (DGE) analyses were performed using either edgeR [69] or the Wilcoxon rank-sum test, depending on the analytical objective. Analyses were restricted to genes present in all sample-level expression matrices and differential expression was defined using a nominal *P*-value < 0.05. Marker genes used to define transcriptional cell populations were identified using the Wilcoxon rank-sum test implemented in Scanpy (v1.11.0) [66], comparing each cluster with the relevant remaining cell populations. EdgeR was used for DGE analysis of S versus D EEC among epithelial cells.

To identify disease-associated transcriptional differences independently of changes in cell abundance, within-cluster DGE analyses were performed using edgeR implemented in the Libra (v1.0.0) [70] R package. A cluster-comparison combination was analysed only when more than 50 cells were available from each diagnostic group. Within each cluster, gene expression values from cells belonging to the same biological sample were summed to generate sample-level pseudobulk profiles, treating samples from the same diagnostic group as replicates.

Complete DGE results for immune and epithelial cells are provided in **Supplementary Table 3** and **Supplementary Table 4** respectively.

### Gene set enrichment analysis

Gene set enrichment analysis (GSEA) was performed on within-clustere edgeR comparisons between disease states using the fgsea (v1.32.4) R package [71]. Genes were ranked according to their average log-fold change derived from the edgeR outputs. Gene sets were obtained from MSigDB (v2024)[72] and included Hallmark pathways, C2 Reactome canonical pathways, and C5 Gene Ontology biological process and molecular functions collections. The C3 transcription factor target collection was additionally included for the epithelial cell analysis.

Where applicable, customised gene signatures derived from previously published immune cell states and manually curated epithelial signals were additionally utilised **(Supplementary Table 5)**.

Pathways passing an FDR threshold of <0.05 were considered significantly enriched. Where no pathway remained significant after multiple-testing correction, directional enrichment was reported only where concordant with gene-level changes and was explicitly described as non-significant. Some of the signatures were renamed before visualisation.

### SCENIC analysis of epithelial regulatory programmes

Gene-regulatory network activity was inferred using pyScenic (0.12.1) [73]. SCENIC analyses were performed independently for ACD and RCD1 epithelial cells. Transcription factor-target relationships were inferred from gene co-expression and refined using transcription-factor motif enrichment, and regulon activity was quantified at single-cell resolution using AUCell.

For each epithelial population, regulons were ranked according to their relative activity after **scaling each regulon to its median activity across all epithelial clusters**. This approach prioritised regulons showing selective activity within individual epithelial populations rather than regulons exhibiting uniformly high activity across the epithelial compartment.

The highest-ranking regulons were examined together with their predicted target genes and interpreted alongside differential-expression and pathway-enrichment analyses.

### Spatial transcriptomics

Formalin-fixed paraffin-embedded (FFPE) duodenal biopsy specimens were sectioned at the Westmead Institute for Medical Research. RNA quality was assessed from representative sections by DV200, with samples with a DV200 >50% taken forward for spatial transcriptomic analysis. For each sample, a 5 µm section was mounted onto a Visium Spatial Gene Expression slide (10x Genomics), deparaffinised and stained with haematoxylin and eosin (H&E). Sections were imaged at 40× magnification using an Olympus whole-slide scanner, and tissue morphology and histopathological features were reviewed by gastrointestinal pathologists with expertise in coeliac disease.Spatial gene-expression profiling was performed using the **Visium Spatial Gene Expression for FFPE assay** with the **Visium Human Transcriptome Probe Set v1.0** (10x Genomics), according to the manufacturer’s instructions. Following imaging and tissue decrosslinking, transcript-specific probe pairs were hybridised and ligated, and spatially barcoded sequencing libraries were generated from the captured ligation products.Final libraries were assessed for fragment-size distribution using an Agilent TapeStation, pooled and sequenced on an Illumina NovaSeq platform to a target depth of approximately 300 million reads per sample.

### Spatial transcriptomics analysis

Spatial transcriptomic profiling was performed on duodenal biopsy sections using the 10x Genomics Visium platform. Sequencing data were processed using the 10x Genomics Space Ranger pipeline (v4.0.1), including alignment to the human reference genome (GRCh38-2020-A), assignment of sequencing reads to spatial barcodes and generation of spot-level gene expression matrices. Tissue-associated spots were identified using the corresponding histological images. Downstream spatial transcriptomic analyses were performed in Python (v3.12.7). Gene expression matrices were normalised using Scanpy (v1.11.0).

Regions of interests (ROIs) were defined by integrating tissue morphology with spatial expression of established epithelial and immune lineage markers. ROIs were selected to represent anatomically and biologically distinct mucosal microenvironments, including villus/surface epithelium, crypt-rich and enteroendocrine-enriched regions, immune-infiltrated epithelial regions, lymphoid aggregates and, where present, glandular regions. Histological annotation was guided by the corresponding tissue image, and markers for each subset of cells were taken from Fitrzpatrick *et al.* [24].

ROI boundaries were subsequently mapped onto the spatial transcriptomic data and all tissue-covered Visium spots within each ROI were extracted for downstream analysis.

For each ROI, expression was aggregated across constituent spots to characterise the relative representation of epithelial and immune transcriptional programmes. Given the multicellular resolution of Visium, spatial co-occurrence was interpreted as evidence of shared or adjacent tissue microenvironments rather than direct cell–cell interaction.

### Data visualisation

Figures were generated in either R (v4.5.2) or Python (3.12.7). In R, the ggplot2 (v4.0.2) package [74] was mainly used for generating plots. TCR gene pairing relationships were visualized as circular chord diagrams using the circlize (v0.4.18) package [75], and the gene usage flows were depicted as alluvial diagrams using the ggaluvial (v0.12.6) package [76] in R. Volcano plots from DGE results were generated using the matplotlib (v3.10.9) python package. Expression of representative genes and predefined signatures were visualised using dot plots (package/software). Spatial expression of individual genes and transcriptional programmes identified by scRNA-seq were visualised across each tissue section using python package xxx.

The schematic representation of the study plan in **Fig. 1A** was generated using BioRender.com.

## Data availability statement

All data associated with this study are present in the paper or the Supplementary Materials. Single-cell gene and protein expression data are available from the National Center for Biotechnology Information (NCBI) under Bioproject PRJNAXXX Custom scripts and data required to perform the analyses presented in this study were deposited in Zenodo at the link: <u>XXXX</u>.

## Ethics statements

### Patient consent for publication

Consent obtained directly from patients.

### Ethics approval

The study was approved by the relevant Human Research Ethics Committees, including Melbourne Health (2020.162) and Western Sydney Local Health District (2021/ETH01429). All participants provided informed consent.

## Acknowledgments

We are grateful to all the volunteers who participated in this study.

## Author contributions

M.S. and F.L. conceived and supervised the study and designed the experiments, with intellectual input from C.C.G. M.S., T.A., S.R., V.V. and J.X. processed clinical samples, performed flow cytometry and contributed to the generation of the single-cell genomic datasets. M.L. and A.S. performed the principal computational and bioinformatic analyses, with major contributions from J.S. and R.L. E.R.B., M.B., B.G., K.J., A.C. and M.F. contributed to data processing, computational analysis and interpretation. J.A.T.-D., G.A., R.C., M.V.L., A.D.S., L.E. and D.S. contributed clinical specimens, clinical data and associated clinical expertise. M.L., A.S., M.S. and F.L. interpreted the data and drafted the manuscript, with input from all authors. All authors contributed to discussion of the results, reviewed the manuscript and approved the final version.

## Supplementary Tables and Figures

**Supplementary Table 1:** Patient metadata

**Supplementary Table 2:** CITE-seq panel

**Supplementary Table 3:** DGE of immune cells

Sheet 1, DGE for cluster genes

Sheet 2, DGE edgeR within cluster comparison Healthy vs ACD

Sheet 3, DGE edgeR within cluster comparison ACD vs RCD1

Sheet 4, DGE between large TCRαβ and Aberrant αβ clones GZMK^+^

Sheet 5, DGE between large TCRαβ and Aberrant αβ clones T effector

**Supplementary Table 4:** DGE of epithelial cells

Sheet 1, DGE within cluster comparison Healthy vs ACD

Sheet 2, DGE within cluster comparison ACD vs RCD1

Sheet 3, DGE EEC + EEC progenitor Healthy vs ACD

Sheet 4, DGE EEC + EEC progenitor ACD vs RCD1

Sheet 5, DGE between EEC D vs S subtype

**Supplementary Table 5:** Customised gene signatures for immnue and epithelial cell subsets

**Supplementary Figure 1. Flow cytometric isolation, sample composition and annotation of duodenal immune cell states.**

**A)** Representative flow-cytometry gating strategy used to isolate live single cells and separate CD45^+^ immune and EPCAM^+^ epithelial populations from dissociated duodenal biopsies. **B)** Stacked bar plots show the relative composition of transcriptionally defined immune-cell populations in Healthy, ACD, RCD1 and RCD2 samples. Colours indicate individual immune cell clusters. **C)** UMAP representations of the immune cell atlas coloured by diagnosis (top) and individual sample identity (bottom), illustrating integration of cells across disease groups and donors. **D)** UMAP representations show mean expression of selected markers used to support immune cell annotation, including CITE-seq surface proteins and selected transcriptional markers. **E)** The dot plot shows expression of selected lineage- and cluster-defining genes across IEL clusters. Dot size represents the proportion of cells expressing each gene and colour represents mean log-transformed normalised expression scaled independently for each geneACD, active coeliac disease; IEL, intraepithelial lymphocyte; ILC, innate lymphoid cell; RCD1 and RCD2, refractory coeliac disease types 1 and 2; UMAP, uniform manifold approximation and projection.

**Supplementary Figure 2. TCR repertoire diversity, clonality and inter-cluster sharing across intestinal T-cell populations.**

**A)** Normalised Shannon entropy of the TCRαβ repertoire across IEL populations (top) and the TCRγδ repertoire across γδ-containing IEL populations (bottom) in Healthy, ACD and RCD1 samples. Higher values indicate greater repertoire diversity. **B)** Sample-level frequencies of singleton TCRγδ clones (right) across Healthy, ACD and RCD1. Each point represents one donor. Displayed p-values are raw two-sided Wilcoxon rank-sum p-values without correction for multiple testing. **C)** Donor-level frequencies of singleton TCRαβ cells (left) and cells belonging to large TCRαβ clones (right) across pooled non-IEL CD4 T-cell and ILC populations in Healthy, ACD and RCD1. Each point represents one donor. Displayed p-values are raw two-sided Wilcoxon rank-sum p-values without correction for multiple testing. **D)** Stacked bar plots showing TCRαβ clone-size distributions across non-IEL CD4 T-cell and ILC populations in Healthy, ACD and RCD1. Only cells containing paired TCRαβ CDR3 sequences are included. Clone-size categories correspond to those defined for the TCRαβ repertoire in Fig. 2. Values in parentheses indicate the number of eligible cells in each population and diagnostic group. **E)** UpSet plot showing shared TCRαβ clonotypes across transcriptionally defined T-cell populations. Connected dots indicate the populations contributing to each clonotype intersection; vertical bars show the number of shared clonotypes for each intersection and horizontal bars show the total number of shared clonotypes associated with each population. Colours indicate diagnostic group. **F)** Corresponding UpSet analysis of shared TCRγδ clonotypes across γδ-containing IEL populations. Connected dots indicate the populations contributing to each clonotype intersection, with bar colours indicating diagnostic group. **G)** Sample-level frequency of *TRDV1–TRGV4* pairing among cells containing paired TCRγδ sequences in Healthy, ACD and RCD1 samples. Each point represents one donor; displayed p-values are raw two-sided Wilcoxon rank-sum p-values.

ACD, active coeliac disease; CDR3, complementarity-determining region 3; IEL, intraepithelial lymphocyte; ILC, innate lymphoid cell; RCD1, refractory coeliac disease type 1; TCR, T-cell receptor.

**Supplementary Figure 3. *GZMK*-high IEL populations show shared and disease-associated transcriptional remodelling.**

**A)** Boxplots show sample-level mean *ITGAE* and *CD69* gene expressions and sample-level mean normalised CD69 and CD57 protein expression across IEL clusters. Each point represents one sample-cluster summary. Boxes show the median and interquartile range, and whiskers extend to 1.5 times the interquartile range. **B)** Dot plots represent selected genes from cluster-specific DGE analysis of four GZMK-enriched clusters. DGE analysis was performed between healthy and ACD, and between ACD and RCD1 using pseudobulk edgeR analysis. Blank positions indicate that the gene was not present in the corresponding DGE result or that the comparison was unavailable because of insufficient cells or samples. **C)** Dot plots show normalised expression of the same genes across disease states. Mean normalised expression and the percentage of expressing cells were first calculated separately for each patient, cluster and gene. Patient-level mean values were then averaged within each diagnostic group, giving each patient equal weight. Dot size represents the patient-balanced mean percentage of cells expressing the gene. Patient-balanced mean expression values were log-transformed using log1p and standardised (z-score) separately for each gene across diagnostic groups. Dot colour therefore represents relative expression of each gene across disease states rather than absolute expression differences between genes.

ACD, active coeliac disease; IEL, intraepithelial lymphocyte; RCD1 and RCD2, refractory coeliac disease types 1 and 2.

**Supplementary Figure 4. SCENIC identifies distinct epithelial regulatory programmes in ACD and RCD1.**

**A–B)** SCENIC analysis of epithelial cells from ACD (**A**) and RCD1 (**B**), performed independently for each disease group. Heatmaps show relative activity of transcription-factor regulons across transcriptionally defined epithelial populations. Regulon activity was quantified using AUCell and ranked according to relative activity after scaling each regulon to its median activity across epithelial clusters, thereby prioritising regulons selectively active within individual epithelial states. Distinct regulatory architectures were observed across the transit-amplifying compartment, including disease-associated differences in Activated TA and Absorptive TA REG3A^+^ populations.

ACD, active coeliac disease; AUCell, area under the recovery curve for regulon activity; RCD1, refractory coeliac disease type 1; SCENIC, single-cell regulatory network inference and clustering; TA, transit-amplifying.

**Supplementary Figure 5. Spatial transcriptomics of additional RCD1, Healthy and ACD duodenal sections support epithelial remodelling and focal epithelial–immune microenvironments.**

**A)** H&E-stained section from an additional RCD1 biopsy (RCD1-16) showing three histology- and marker-guided regions of interest (ROIs). **B)** Spatial expression of selected genes across the RCD1 section. Colour represents log-transformed normalised gene expression. **C)** The dot plot shows selected epithelial and immune programmes across the three RCD1 ROIs. Genes are grouped and coloured according to functional and structural programmes. Dot size represents the percentage of Visium spots within each ROI expressing the indicated gene, and colour represents scaled mean log-transformed normalised expression across spots within each ROI. **D)** H&E-stained duodenal section from a Healthy control (H-9). **E)** Spatial expression of selected genes in the Healthy section. Colour represents log-transformed normalised gene expression. **F)** H&E-stained tissue from the first section of an ACD biopsy (ACD-1). **G)** Spatial expression of the same genes in **E** across the first ACD section. **H)** H&E-stained tissue from a second section of the same ACD biopsy (ACD-1). **I)** Spatial expression of the same genes in **E** across the second ACD section. H&E, haematoxylin and eosin; ACD, active coeliac disease; EEC, enteroendocrine; RCD1, refractory coeliac disease type 1; ROI, region of interest.

