## Supplementary Figures for "Single-cell multi-omics maps clonal IEL expansion and epithelial remodelling in refractory coeliac disease"

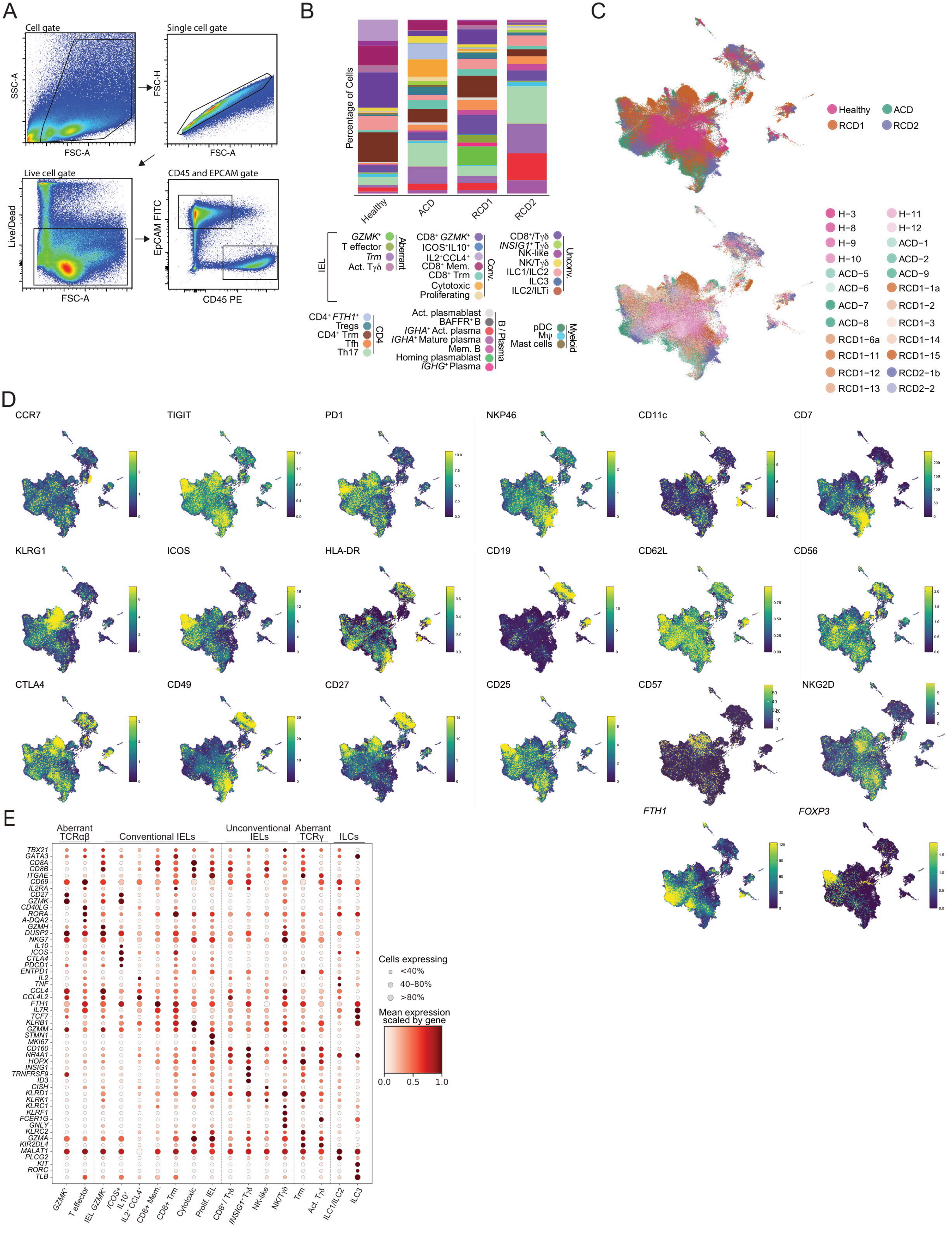
A
Cell gate
SSC-A
FSC-A
Single cell gate
FSC-H
FSC-A
Live cell gate
Live/Dead
FSC-A
CD45 and EPCAM gate
EpCAM FITC
CD45 PE
B
Percentage of Cells
Healthy
ACD
RCD1
RCD2
IEL
Aberrant
GZMK+
T effector
Trm
Act. Tγδ
Conv.
CD8+ GZMK+
ICOS+IL10+
IL2+CCL4+
CD8+ Mem.
CD8+ Trm
Cytotoxic
Proliferating
Unconv.
CD8+/Tγδ
INSIG1+ Tγδ
NK-like
NK/Tγδ
ILC1/ILC2
ILC3
ILC2/ILTi
CD4
CD4+ FTH1+
Tregs
CD4+ Trm
Tfh
Th17
B / Plasma
Act. plasmablast
BAFFR+ B
IGHA+ Act. plasma
IGHA+ Mature plasma
Mem. B
Homing plasmablast
IGHG+ Plasma
Myeloid
pDC
Mψ
Mast cells
C
Healthy
ACD
RCD1
RCD2
H−3
H−8
H−9
H−10
ACD−5
ACD−6
ACD−7
ACD−8
RCD1−6a
RCD1−11
RCD1−12
RCD1−13
H−11
H−12
ACD−1
ACD−2
ACD−9
RCD1−1a
RCD1−2
RCD1−3
RCD1−14
RCD1−15
RCD2−1b
RCD2−2
D
CCR7
TIGIT
PD1
NKP46
CD11c
CD7
KLRG1
ICOS
HLA-DR
CD19
CD62L
CD56
CTLA4
CD49
CD27
CD25
CD57
NKG2D
FTH1
FOXP3
E
Aberrant TCRαβ
Conventional IELs
Unconventional IELs
Aberrant TCRγ
ILCs
TBX21
GATA3
CD8A
CD8B
ITGAE
CD69
IL2RA
CD27
GZMK
CD40LG
RORA
A-DQA2
GZMH
DUSP2
NKG7
IL10
ICOS
CTLA4
PDCD1
ENTPD1
IL2
TNF
CCL4
CCL4L2
FTH1
IL7R
KLRB1
TCF7
GZMM
STMN1
MKI67
CD160
NR4A1
HOPX
INSIG1
TRNFRSF9
ID3
CISH
KLRD1
KLRK1
KLRC1
KLRF1
FCER1G
GNLY
KLRC2
GZMA
KIR2DL4
MALAT1
PLCG2
KIT
RORC
TLB
GZMK+
T effector
IEL GZMK+
ICOS+ IL10+
IL2+ CCL4+
CD8+ Mem.
CD8+ Trm
Cytotoxic
Prolif. IEL
CD8+ / Tγδ
INSIG1+ Tγδ
NK-like
NK/Tγδ
Trm
Act. Tγδ
ILC1/ILC2
ILC3
Cells expressing
<40%
40–80%
>80%
Mean expression scaled by gene
0.0
0.5
1.0

**Supplementary Figure 1. Flow cytometric isolation, sample composition and annotation of duodenal immune cell states.**

**A)** Representative flow-cytometry gating strategy used to isolate live single cells and separate $CD45^+$ immune and $EPCAM^+$ epithelial populations from dissociated duodenal biopsies. **B)** Stacked bar plots show the relative composition of transcriptionally defined immune-cell populations in Healthy, ACD, RCD1 and RCD2 samples. Colours indicate individual immune cell clusters. **C)** UMAP representations of the immune cell atlas coloured by diagnosis (top) and individual sample identity (bottom), illustrating integration of cells across disease groups and donors. **D)** UMAP representations show mean expression of selected markers used to support immune cell annotation, including CITE-seq surface proteins and selected transcriptional markers. **E)** The dot plot shows expression of selected lineage- and cluster-defining genes across IEL clusters. Dot size represents the proportion of cells expressing each gene and colour represents mean log-transformed normalised expression scaled independently for each geneACD, active coeliac disease; IEL, intraepithelial lymphocyte; ILC, innate lymphoid cell; RCD1 and RCD2, refractory coeliac disease types 1 and 2; UMAP, uniform manifold approximation and projection.

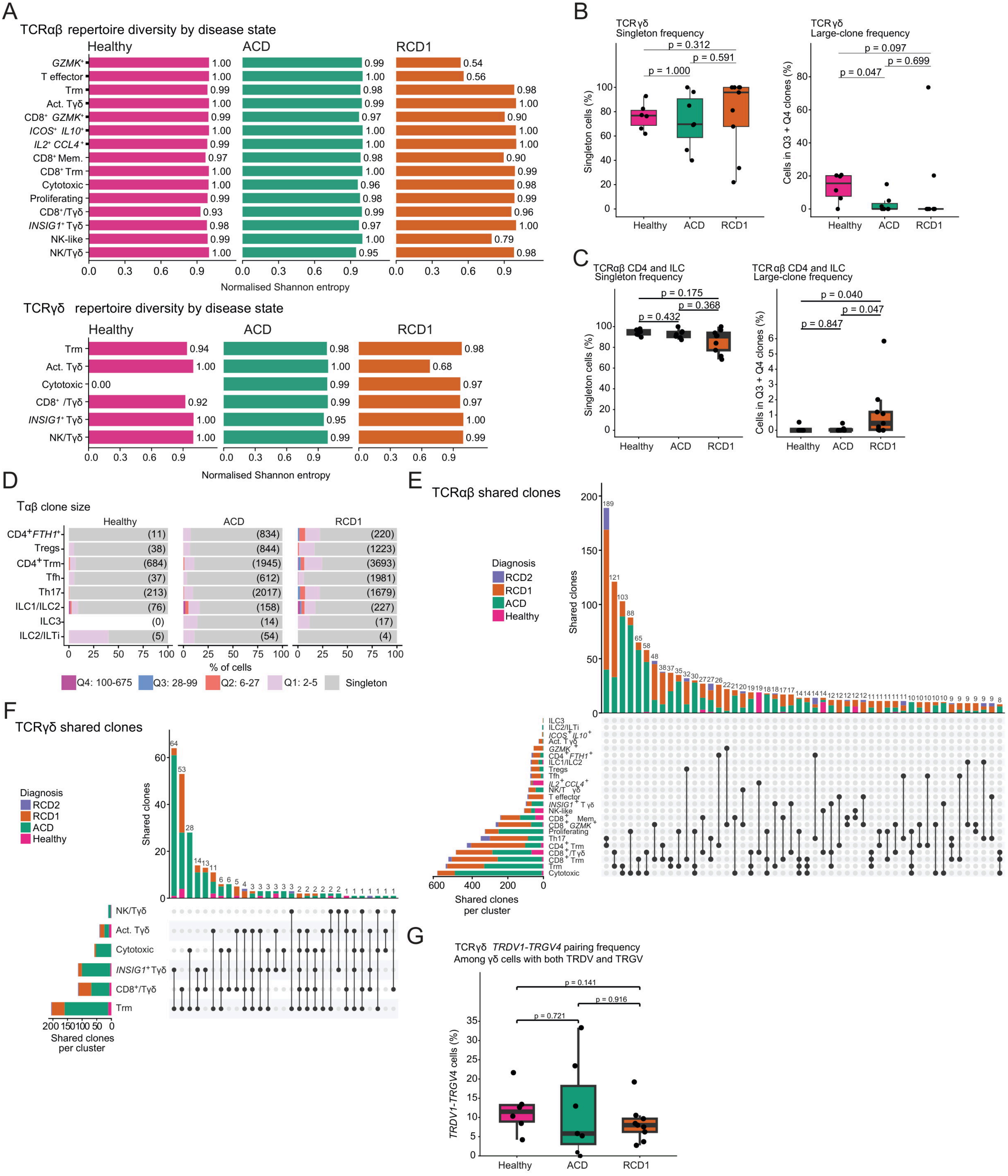
A
TCRαβ repertoire diversity by disease state
Healthy
ACD
RCD1
GZMK+
T effector
Trm
Act. Tγδ
CD8+ GZMK+
ICOS+ IL10+
IL2+ CCL4+
CD8+ Mem.
CD8+ Trm
Cytotoxic
Proliferating
CD8+/Tγδ
INSIG1+ Tγδ
NK-like
NK/Tγδ
Normalised Shannon entropy
TCRγδ repertoire diversity by disease state
Healthy
ACD
RCD1
Trm
Act. Tγδ
Cytotoxic
CD8+ /Tγδ
INSIG1+ Tγδ
NK/Tγδ
Normalised Shannon entropy
B
TCRγδ
Singleton frequency
p = 0.312
p = 0.591
p = 1.000
Singleton cells (%)
Healthy
ACD
RCD1
TCRγδ
Large-clone frequency
p = 0.097
p = 0.699
p = 0.047
Cells in Q3 + Q4 clones (%)
Healthy
ACD
RCD1
C
TCRαβ CD4 and ILC
Singleton frequency
p = 0.175
p = 0.368
p = 0.432
Singleton cells (%)
Healthy
ACD
RCD1
TCRαβ CD4 and ILC
Large-clone frequency
p = 0.040
p = 0.047
p = 0.847
Cells in Q3 + Q4 clones (%)
Healthy
ACD
RCD1
D
Tαβ clone size
Healthy
ACD
RCD1
CD4+FTH1+
Tregs
CD4+Trm
Tfh
Th17
ILC1/ILC2
ILC3
ILC2/ILTi
% of cells
Q4: 100-675
Q3: 28-99
Q2: 6-27
Q1: 2-5
Singleton
E
TCRαβ shared clones
Diagnosis
RCD2
RCD1
ACD
Healthy
Shared clones
Shared clones per cluster
F
TCRγδ shared clones
Diagnosis
RCD2
RCD1
ACD
Healthy
Shared clones
NK/Tγδ
Act. Tγδ
Cytotoxic
INSIG1+Tγδ
CD8+/Tγδ
Trm
Shared clones per cluster
G
TCRγδ TRDV1-TRGV4 pairing frequency
Among γδ cells with both TRDV and TRGV
p = 0.141
p = 0.916
p = 0.721
TRDV1-TRGV4 cells (%)
Healthy
ACD
RCD1

**Supplementary Figure 2. TCR repertoire diversity, clonality and inter-cluster sharing across intestinal T-cell populations.**

**A)** Normalised Shannon entropy of the TCRαβ repertoire across IEL populations (top) and the TCRγδ repertoire across γδ-containing IEL populations (bottom) in Healthy, ACD and RCD1 samples. Higher values indicate greater repertoire diversity. **B)** Sample-level frequencies of singleton TCRγδ clones (right) across Healthy, ACD and RCD1. Each point represents one donor. Displayed p-values are raw two-sided Wilcoxon rank-sum p-values without correction for multiple testing. **C)** Donor-level frequencies of singleton TCRαβ cells (left) and cells belonging to large TCRαβ clones (right) across pooled non-IEL CD4 T-cell and ILC populations in Healthy, ACD and RCD1. Each point represents one donor. Displayed p-values are raw two-sided Wilcoxon rank-sum p-values without correction for multiple testing. **D)** Stacked bar plots showing TCRαβ clone-size distributions across non-IEL CD4 T-cell and ILC populations in Healthy, ACD and RCD1. Only cells containing paired TCRαβ CDR3 sequences are included. Clone-size categories correspond to those defined for the TCRαβ repertoire in Fig. 2. Values in parentheses indicate the number of eligible cells in each population and diagnostic group. **E)** UpSet plot showing shared TCRαβ clonotypes across transcriptionally defined T-cell populations. Connected dots indicate the populations contributing to each clonotype intersection; vertical bars show the number of shared clonotypes for each intersection and horizontal bars show the total number of shared clonotypes associated with each population. Colours indicate diagnostic group. **F)** Corresponding UpSet analysis of shared TCRγδ clonotypes across γδ-containing IEL populations. Connected dots indicate the populations contributing to each clonotype intersection, with bar colours indicating diagnostic group. **G)** Sample-level frequency of *TRDV1–TRGV4* pairing among cells containing paired TCRγδ sequences in Healthy, ACD and RCD1 samples. Each point represents one donor; displayed p-values are raw two-sided Wilcoxon rank-sum p-values. ACD, active coeliac disease; CDR3, complementarity-determining region 3; IEL, intraepithelial lymphocyte; ILC, innate lymphoid cell; RCD1, refractory coeliac disease type 1; TCR, T-cell receptor.

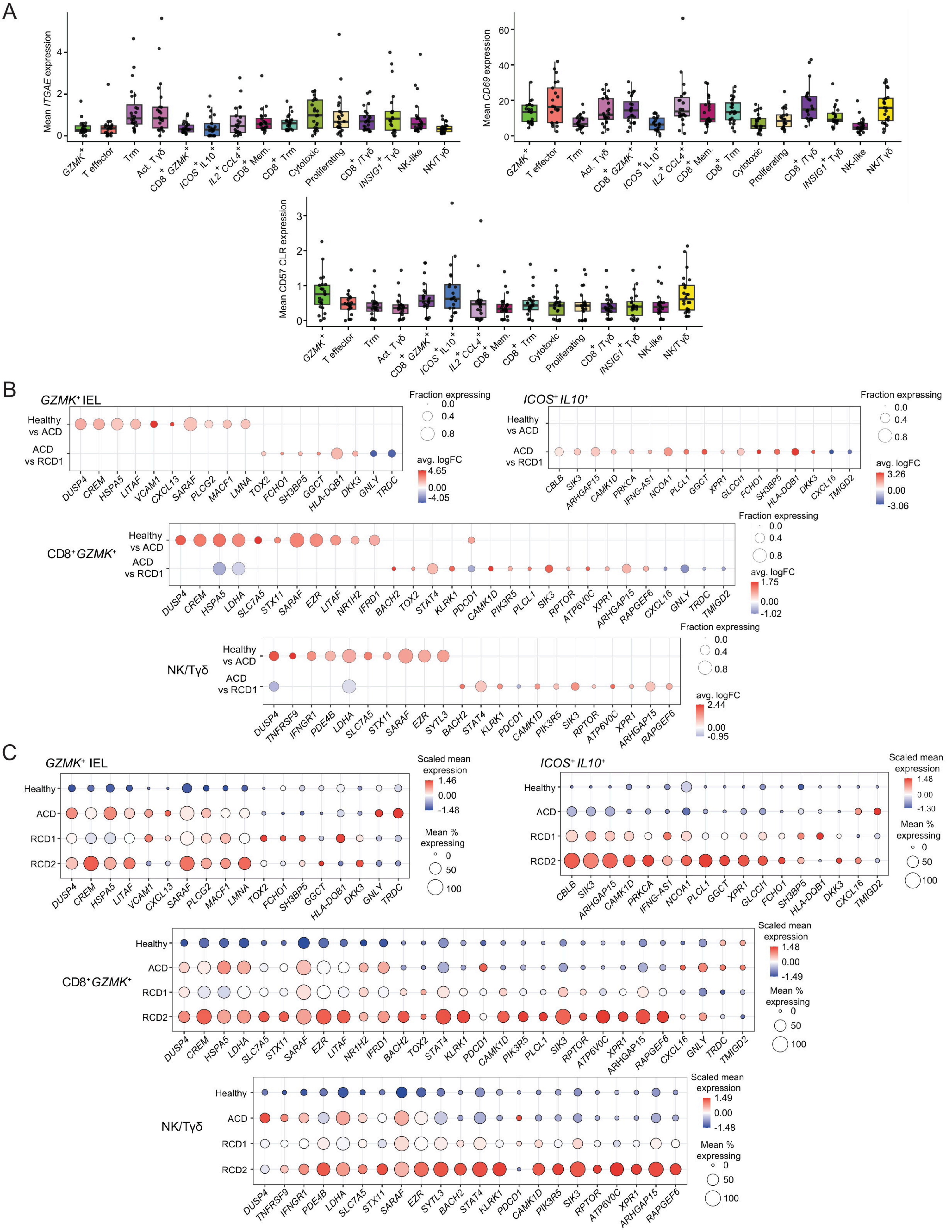
A
Mean ITGAE expression
Mean CD69 expression
Mean CD57 CLR expression
GZMK+
T effector
Trm
Act. Tγδ
CD8+ GZMK+
ICOS+ IL10+
IL2+ CCL4+
CD8+ Mem.
CD8+ Trm
Cytotoxic
Proliferating
CD8+/Tγδ
INSIG1+ Tγδ
NK-like
NK/Tγδ
B
GZMK+ IEL
ICOS+ IL10+
CD8+ GZMK+
NK/Tγδ
Healthy vs ACD
ACD vs RCD1
Fraction expressing
avg. logFC
DUSP4
CREM
HSPA5
LITAF
VCAM1
CXCL13
SARAF
PLCG2
MACF1
LMNA
TOX2
FCHO1
SH3BP5
GGCT
HLA-DQB1
DKK3
GNLY
TRDC
CBLB
SIK3
ARHGAP15
CAMK1D
PRKCA
IFNG-AS1
NCOA1
PLCL1
XPR1
GLCCI1
CXCL16
TMIGD2
LDHA
SLC7A5
STX11
EZR
NR1H2
IFRD1
BACH2
STAT4
KLRK1
PDCD1
PIK3R5
RPTOR
ATP6V0C
RAPGEF6
TNFRSF9
IFNGR1
PDE4B
SYTL3
C
Healthy
ACD
RCD1
RCD2
Scaled mean expression
Mean % expressing

**Supplementary Figure 3.** ***GZMK*****-high IEL populations show shared and disease-associated transcriptional remodelling.**

**A)** Boxplots show sample-level mean *ITGAE* and *CD69* gene expressions and sample-level mean normalised CD69 and CD57 protein expression across IEL clusters. Each point represents one sample-cluster summary. Boxes show the median and interquartile range, and whiskers extend to 1.5 times the interquartile range. **B)** Dot plots represent selected genes from cluster-specific DGE analysis of four GZMK-enriched clusters. DGE analysis was performed between healthy and ACD, and between ACD and RCD1 using pseudobulk edgeR analysis. Blank positions indicate that the gene was not present in the corresponding DGE result or that the comparison was unavailable because of insufficient cells or samples. **C)** Dot plots show normalised expression of the same genes across disease states. Mean normalised expression and the percentage of expressing cells were first calculated separately for each patient, cluster and gene. Patient-level mean values were then averaged within each diagnostic group, giving each patient equal weight. Dot size represents the patient-balanced mean percentage of cells expressing the gene. Patient-balanced mean expression values were log-transformed using log1p and standardised (z-score) separately for each gene across diagnostic groups. Dot colour therefore represents relative expression of each gene across disease states rather than absolute expression differences between genes.

ACD, active coeliac disease; IEL, intraepithelial lymphocyte; RCD1 and RCD2, refractory coeliac disease types 1 and 2.

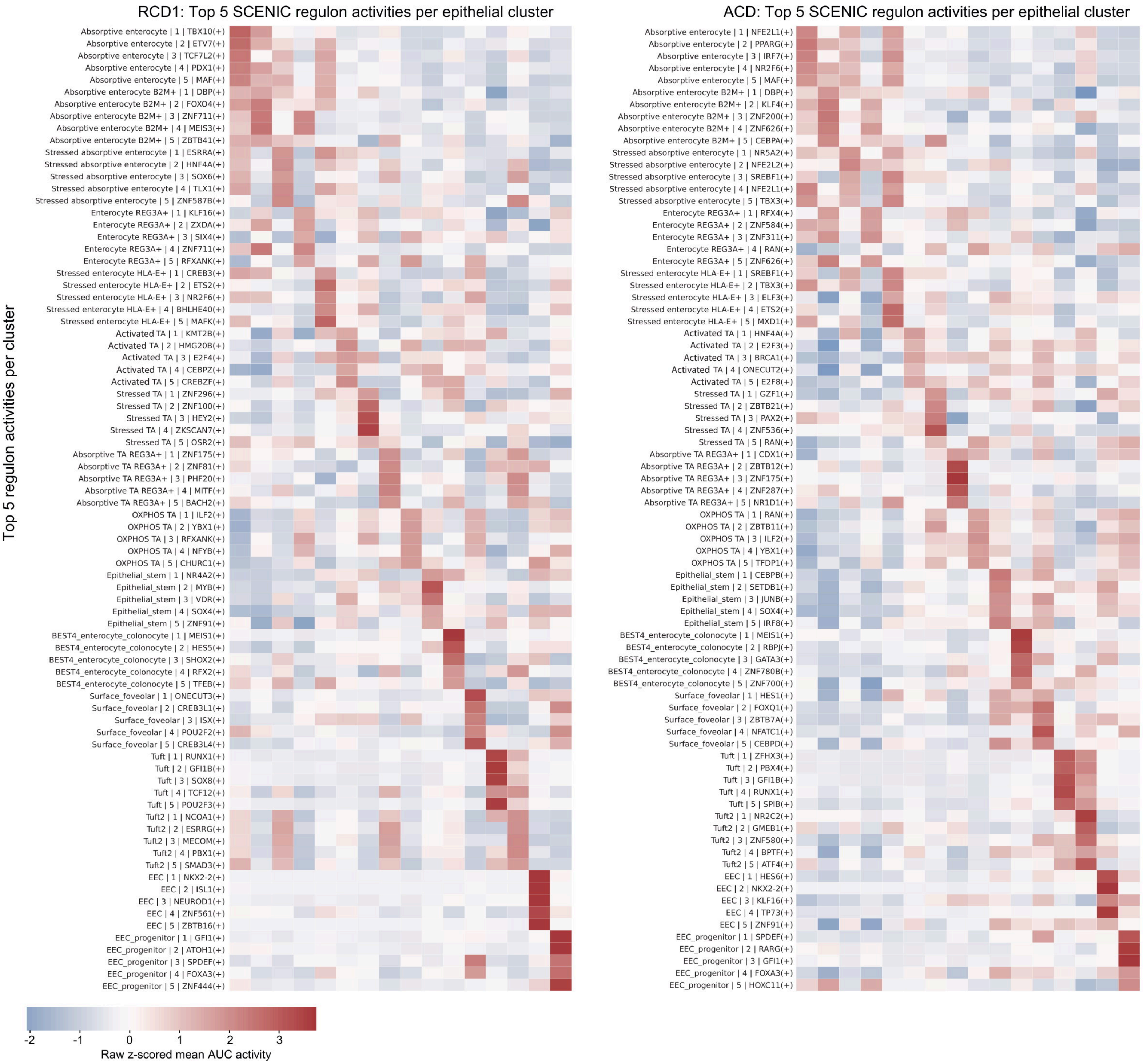
RCD1: Top 5 SCENIC regulon activities per epithelial cluster
Top 5 regulon activities per cluster
Absorptive enterocyte | 1 | TBX10(+)
Absorptive enterocyte | 2 | ETV7(+)
Absorptive enterocyte | 3 | TCF7L2(+)
Absorptive enterocyte | 4 | PDX1(+)
Absorptive enterocyte | 5 | MAF(+)
Absorptive enterocyte B2M+ | 1 | DBP(+)
Absorptive enterocyte B2M+ | 2 | FOXO4(+)
Absorptive enterocyte B2M+ | 3 | ZNF711(+)
Absorptive enterocyte B2M+ | 4 | MEIS3(+)
Absorptive enterocyte B2M+ | 5 | ZBTB41(+)
Stressed absorptive enterocyte | 1 | ESRRA(+)
Stressed absorptive enterocyte | 2 | HNF4A(+)
Stressed absorptive enterocyte | 3 | SOX6(+)
Stressed absorptive enterocyte | 4 | TLX1(+)
Stressed absorptive enterocyte | 5 | ZNF587B(+)
Enterocyte REG3A+ | 1 | KLF16(+)
Enterocyte REG3A+ | 2 | ZXDA(+)
Enterocyte REG3A+ | 3 | SIX4(+)
Enterocyte REG3A+ | 4 | ZNF711(+)
Enterocyte REG3A+ | 5 | RFXANK(+)
Stressed enterocyte HLA-E+ | 1 | CREB3(+)
Stressed enterocyte HLA-E+ | 2 | ETS2(+)
Stressed enterocyte HLA-E+ | 3 | NR2F6(+)
Stressed enterocyte HLA-E+ | 4 | BHLHE40(+)
Stressed enterocyte HLA-E+ | 5 | MAFK(+)
Activated TA | 1 | KMT2B(+)
Activated TA | 2 | HMG20B(+)
Activated TA | 3 | E2F4(+)
Activated TA | 4 | CEBPZ(+)
Activated TA | 5 | CREBZF(+)
Stressed TA | 1 | ZNF296(+)
Stressed TA | 2 | ZNF100(+)
Stressed TA | 3 | HEY2(+)
Stressed TA | 4 | ZKSCAN7(+)
Stressed TA | 5 | OSR2(+)
Absorptive TA REG3A+ | 1 | ZNF175(+)
Absorptive TA REG3A+ | 2 | ZNF81(+)
Absorptive TA REG3A+ | 3 | PHF20(+)
Absorptive TA REG3A+ | 4 | MITF(+)
Absorptive TA REG3A+ | 5 | BACH2(+)
OXPHOS TA | 1 | ILF2(+)
OXPHOS TA | 2 | YBX1(+)
OXPHOS TA | 3 | RFXANK(+)
OXPHOS TA | 4 | NFYB(+)
OXPHOS TA | 5 | CHURC1(+)
Epithelial_stem | 1 | NR4A2(+)
Epithelial_stem | 2 | MYB(+)
Epithelial_stem | 3 | VDR(+)
Epithelial_stem | 4 | SOX4(+)
Epithelial_stem | 5 | ZNF91(+)
BEST4_enterocyte_colonocyte | 1 | MEIS1(+)
BEST4_enterocyte_colonocyte | 2 | HES5(+)
BEST4_enterocyte_colonocyte | 3 | SHOX2(+)
BEST4_enterocyte_colonocyte | 4 | RFX2(+)
BEST4_enterocyte_colonocyte | 5 | TFEB(+)
Surface_foveolar | 1 | ONECUT3(+)
Surface_foveolar | 2 | CREB3L1(+)
Surface_foveolar | 3 | ISX(+)
Surface_foveolar | 4 | POU2F2(+)
Surface_foveolar | 5 | CREB3L4(+)
Tuft | 1 | RUNX1(+)
Tuft | 2 | GFI1B(+)
Tuft | 3 | SOX8(+)
Tuft | 4 | TCF12(+)
Tuft | 5 | POU2F3(+)
Tuft2 | 1 | NCOA1(+)
Tuft2 | 2 | ESRRG(+)
Tuft2 | 3 | MECOM(+)
Tuft2 | 4 | PBX1(+)
Tuft2 | 5 | SMAD3(+)
EEC | 1 | NKX2-2(+)
EEC | 2 | ISL1(+)
EEC | 3 | NEUROD1(+)
EEC | 4 | ZNF561(+)
EEC | 5 | ZBTB16(+)
EEC_progenitor | 1 | GFI1(+)
EEC_progenitor | 2 | ATOH1(+)
EEC_progenitor | 3 | SPDEF(+)
EEC_progenitor | 4 | FOXA3(+)
EEC_progenitor | 5 | ZNF444(+)
ACD: Top 5 SCENIC regulon activities per epithelial cluster
Absorptive enterocyte | 1 | NFE2L1(+)
Absorptive enterocyte | 2 | PPARG(+)
Absorptive enterocyte | 3 | IRF7(+)
Absorptive enterocyte | 4 | NR2F6(+)
Absorptive enterocyte | 5 | MAF(+)
Absorptive enterocyte B2M+ | 1 | DBP(+)
Absorptive enterocyte B2M+ | 2 | KLF4(+)
Absorptive enterocyte B2M+ | 3 | ZNF200(+)
Absorptive enterocyte B2M+ | 4 | ZNF626(+)
Absorptive enterocyte B2M+ | 5 | CEBPA(+)
Stressed absorptive enterocyte | 1 | NR5A2(+)
Stressed absorptive enterocyte | 2 | NFE2L2(+)
Stressed absorptive enterocyte | 3 | SREBF1(+)
Stressed absorptive enterocyte | 4 | NFE2L1(+)
Stressed absorptive enterocyte | 5 | TBX3(+)
Enterocyte REG3A+ | 1 | RFX4(+)
Enterocyte REG3A+ | 2 | ZNF584(+)
Enterocyte REG3A+ | 3 | ZNF311(+)
Enterocyte REG3A+ | 4 | RAN(+)
Enterocyte REG3A+ | 5 | ZNF626(+)
Stressed enterocyte HLA-E+ | 1 | SREBF1(+)
Stressed enterocyte HLA-E+ | 2 | TBX3(+)
Stressed enterocyte HLA-E+ | 3 | ELF3(+)
Stressed enterocyte HLA-E+ | 4 | ETS2(+)
Stressed enterocyte HLA-E+ | 5 | MXD1(+)
Activated TA | 1 | HNF4A(+)
Activated TA | 2 | E2F3(+)
Activated TA | 3 | BRCA1(+)
Activated TA | 4 | ONECUT2(+)
Activated TA | 5 | E2F8(+)
Stressed TA | 1 | GZF1(+)
Stressed TA | 2 | ZBTB21(+)
Stressed TA | 3 | PAX2(+)
Stressed TA | 4 | ZNF536(+)
Stressed TA | 5 | RAN(+)
Absorptive TA REG3A+ | 1 | CDX1(+)
Absorptive TA REG3A+ | 2 | ZBTB12(+)
Absorptive TA REG3A+ | 3 | ZNF175(+)
Absorptive TA REG3A+ | 4 | ZNF287(+)
Absorptive TA REG3A+ | 5 | NR1D1(+)
OXPHOS TA | 1 | RAN(+)
OXPHOS TA | 2 | ZBTB11(+)
OXPHOS TA | 3 | ILF2(+)
OXPHOS TA | 4 | YBX1(+)
OXPHOS TA | 5 | TFDP1(+)
Epithelial_stem | 1 | CEBPB(+)
Epithelial_stem | 2 | SETDB1(+)
Epithelial_stem | 3 | JUNB(+)
Epithelial_stem | 4 | SOX4(+)
Epithelial_stem | 5 | IRF8(+)
BEST4_enterocyte_colonocyte | 1 | MEIS1(+)
BEST4_enterocyte_colonocyte | 2 | RBPJ(+)
BEST4_enterocyte_colonocyte | 3 | GATA3(+)
BEST4_enterocyte_colonocyte | 4 | ZNF780B(+)
BEST4_enterocyte_colonocyte | 5 | ZNF700(+)
Surface_foveolar | 1 | HES1(+)
Surface_foveolar | 2 | FOXQ1(+)
Surface_foveolar | 3 | ZBTB7A(+)
Surface_foveolar | 4 | NFATC1(+)
Surface_foveolar | 5 | CEBPD(+)
Tuft | 1 | ZFHX3(+)
Tuft | 2 | PBX4(+)
Tuft | 3 | GFI1B(+)
Tuft | 4 | RUNX1(+)
Tuft | 5 | SPIB(+)
Tuft2 | 1 | NR2C2(+)
Tuft2 | 2 | GMEB1(+)
Tuft2 | 3 | ZNF580(+)
Tuft2 | 4 | BPTF(+)
Tuft2 | 5 | ATF4(+)
EEC | 1 | HES6(+)
EEC | 2 | NKX2-2(+)
EEC | 3 | KLF16(+)
EEC | 4 | TP73(+)
EEC | 5 | ZNF91(+)
EEC_progenitor | 1 | SPDEF(+)
EEC_progenitor | 2 | RARG(+)
EEC_progenitor | 3 | GFI1(+)
EEC_progenitor | 4 | FOXA3(+)
EEC_progenitor | 5 | HOXC11(+)
-2
-1
0
1
2
3
Raw z-scored mean AUC activity

**Supplementary Figure 4. SCENIC identifies distinct epithelial regulatory programmes in ACD and RCD1.**

**A–B)** SCENIC analysis of epithelial cells from ACD (**A**) and RCD1 (**B**), performed independently for each disease group. Heatmaps show relative activity of transcription-factor regulons across transcriptionally defined epithelial populations. Regulon activity was quantified using AUCell and ranked according to relative activity after scaling each regulon to its median activity across epithelial clusters, thereby prioritising regulons selectively active within individual epithelial states. Distinct regulatory architectures were observed across the transit-amplifying compartment, including disease-associated differences in Activated TA and Absorptive TA REG3A$^{+}$ populations. ACD, active coeliac disease; AUCell, area under the recovery curve for regulon activity; RCD1, refractory coeliac disease type 1; SCENIC, single-cell regulatory network inference and clustering; TA, transit-amplifying.

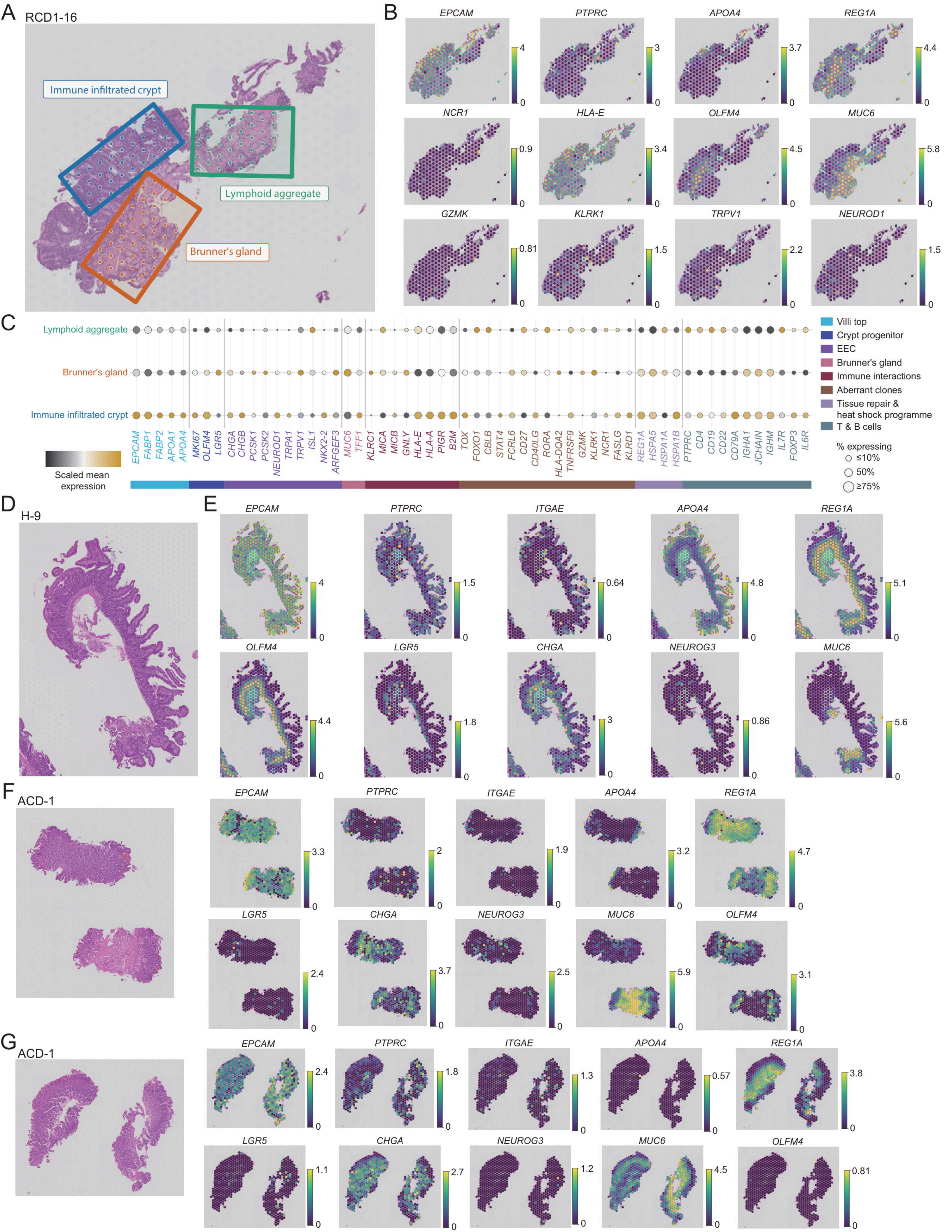

A
RCD1-16
Immune infiltrated crypt
Lymphoid aggregate
Brunner's gland
B
EPCAM
PTPRC
APOA4
REG1A
NCR1
HLA-E
OLFM4
MUC6
GZMK
KLRK1
TRPV1
NEUROD1
C
Lymphoid aggregate
Brunner's gland
Immune infiltrated crypt
Scaled mean expression
Villi top
Crypt progenitor
EEC
Brunner's gland
Immune interactions
Aberrant clones
Tissue repair & heat shock programme
T & B cells
% expressing
≤10%
50%
≥75%
D
H-9
E
ITGAE
LGR5
CHGA
NEUROG3
F
ACD-1
G
ACD-1

**Supplementary Figure 5. Spatial transcriptomics of additional RCD1, Healthy and ACD duodenal sections support epithelial remodelling and focal epithelial–immune microenvironments.**

**A)** H&E-stained section from an additional RCD1 biopsy (RCD1-16) showing three histology- and marker-guided regions of interest (ROIs). **B)** Spatial expression of selected genes across the RCD1 section. Colour represents log-transformed normalised gene expression. **C)** The dot plot shows selected epithelial and immune programmes across the three RCD1 ROIs. Genes are grouped and coloured according to functional and structural programmes. Dot size represents the percentage of Visium spots within each ROI expressing the indicated gene, and colour represents scaled mean log-transformed normalised expression across spots within each ROI. **D)** H&E-stained duodenal section from a Healthy control (H-9). **E)** Spatial expression of selected genes in the Healthy section. Colour represents log-transformed normalised gene expression. **F)** H&E-stained tissue from the first section of an ACD biopsy (ACD-1). **G)** Spatial expression of the same genes in **E** across the first ACD section. **H)** H&E-stained tissue from a second section of the same ACD biopsy (ACD-1). **I)** Spatial expression of the same genes in **E** across the second ACD section. H&E, haematoxylin and eosin; ACD, active coeliac disease; EEC, enteroendocrine; RCD1, refractory coeliac disease type 1; ROI, region of interest.
